# MTHFD2 is a Metabolic Checkpoint Controlling Effector and Regulatory T Cell Fate and Function

**DOI:** 10.1101/2021.02.03.428939

**Authors:** Ayaka Sugiura, Gabriela Andrejeva, Kelsey Voss, Darren R. Heintzman, Katherine L. Beier, Melissa M. Wolf, Dalton Greenwood, Xiang Ye, Shailesh K. Shahi, Samantha N. Freedman, Alanna M. Cameron, Patrik Foerch, Tim Bourne, Xincheng Xu, Juan C. Garcia-Canaveras, Ashutosh K. Mangalam, Joshua D. Rabinowitz, Jeffrey C. Rathmell

## Abstract

Antigenic stimulation promotes T cells metabolic reprogramming to meet increased biosynthetic, bioenergetic, and signaling demands. We show that the one-carbon (1C) metabolism enzyme Methylenetetrahydrofolate Dehydrogenase-2 (MTHFD2) is highly expressed in inflammatory diseases and induced in activated T cells to promote proliferation and produce inflammatory cytokines. In pathogenic Th17 cells, MTHFD2 also prevented aberrant upregulation of FoxP3 and suppressive capacity. Conversely, MTHFD2-deficiency enhanced lineage stability of regulatory T (Treg) cells. Mechanistically, MTHFD2 maintained cellular 10-formyltetrahydrofolate for *de novo* purine synthesis and MTHFD2 inhibition led to accumulation of the intermediate 5-aminoimidazole carboxamide ribonucleotide that was associated with decreased mTORC1 signaling. MTHFD2 was also required for proper histone de-methylation in Th17 cells. Importantly, inhibiting MTHFD2 *in vivo* reduced disease severity in Experimental Autoimmune Encephalomyelitis and Delayed-Type Hypersensitivity. MTHFD2 induction is thus a metabolic checkpoint for pathogenic effector cells that suppresses anti-inflammatory Treg cells and is a potential therapeutic target within 1C metabolism.

## INTRODUCTION

Effective mobilization of the adaptive immune response requires robust activation of CD4 T cells. Upon antigenic stimulation, CD4 T cells undergo rapid cell growth and proliferation. This involves metabolic reprogramming driven by mTORC1 from a catabolic resting state to an anabolic growth state, with increased biosynthetic, bioenergetic, and signaling demands. Depending on the cytokine milieu, CD4 T cells differentiate into effector and regulatory subsets that have distinct metabolic programs driving their effector functions (Bantug et al., 2018; Buck et al., 2015). The balance of these subsets is critical to allow for normal immunity while preventing inflammatory and autoimmune diseases. For example, Multiple Sclerosis (MS) is characterized by increased IL-17-producing Th17 cells and decreased or ineffectual suppressive regulatory T (Treg) cells (Dendrou et al., 2015). Targeting the specific metabolic programs of these T cell subsets to re-establish the appropriate balance offers a new approach to cell metabolism-based immunotherapy.

Cell-metabolism based therapeutics date back to 1948, when Sidney Farber first showed the efficacy of an anti-folate chemotherapeutic in children with acute lymphoblastic leukemia (Farber et al., 1948). Many different chemotherapeutics have since been developed to target this pathway including methotrexate and other anti-folates, fluorouracil (5-FU), and mercaptopurine (6-MP). Broadly, these compounds target the one-carbon (1C) metabolism pathways and nucleotide synthesis. Rapidly proliferating cells, including cancer cells and activated T cells, share dependencies on these pathways to synthesize DNA and RNA. Methotrexate remains commonly used to treat autoimmune and inflammatory diseases, such as rheumatoid arthritis (Brown et al., 2016). However, due to the broad expression of the targeted enzymes, these therapeutics are associated with common and potentially severe adverse effects. Identifying metabolic enzyme targets that are uniquely important in cell populations of interest could lead to the development of safer and more efficacious immunotherapies.

1C metabolism includes the folate and methionine cycles and involves the transfer of single carbon units. These carbon units can then be used for the biosynthesis of purines and thymidylates, generation of methyl donors, as well as maintenance of cellular redox balance (Ducker and Rabinowitz, 2017; Yang and Vousden, 2016). Endogenously synthesized or imported serine serves as a 1C donor to activate tetrahydrofolate (THF) to 5,10-methyleneTHF, with concomitant glycine production. 5,10-methyleneTHF can then be oxidized using NAD(P) to generate the purine precursor 10-formylTHF. Alternatively, 10-formylTHF can be synthesized from formate and THF in an ATP-dependent manner. In the cytosolic pathway, 10-fomylTHF production is mediated by Methylenetetrahydrofolate Dehydrogenase 1 (MTHFD1). The mitochondrial pathway relies on mitochondrial Methylenetetrahydrofolate Dehydrogenase (MTHFD2) and MTHFD1-Like (MTHFD1L). Loss of MTHFD1 completely starves cells for cytosolic 10-formylTHF, ablating purine biosynthetic capacity. Individuals with mutations in MTHFD1 develop Severe Combined Immunodeficiency (SCID) (Field et al., 2015).

MTHFD2 has garnered interest in cancer biology as one of the most highly induced and overexpressed genes in all tumors (Nilsson et al., 2014). Importantly, while MTHFD2 is broadly upregulated during embryogenesis, it has very little to no expression in most adult tissues (Nilsson et al., 2014). As a novel target for anti-cancer therapy (Zhu and Leung, 2020), proposed mechanisms include increased oxidative stress (Ju et al., 2019; Wan et al., 2020), exogenous glycine dependency (Koufaris et al., 2016), and insufficient purine synthesis (Ben-Sahra et al., 2016; Pikman et al., 2016). Additionally, MTHFD2 deficiency can lead to accumulation of the purine synthesis pathway intermediate and adenosine monophosphate (AMP) analog, 5-aminoimidazole carboxamide ribonucleotide (AICAR) (Ducker et al., 2016). AICAR in turn can inhibit cancer cell growth through AMP activated Protein Kinase (AMPK)/mTOR-dependent (Su et al., 2019) and -independent (Dembitz et al., 2019) pathways.

Because 1C metabolism integrates multiple nutrient inputs and is therapeutically tractable, we examined the role of 1C metabolism in CD4 T effector (Teff) and Treg cell subsets. Through unbiased *in vivo* CRISPR-Cas9 based targeted screening in primary murine T cells, we identified MTHFD2 as a hit that was also consistently upregulated in patients with inflammatory disorders. Notably, MTHFD2 deficiency impaired Teff function and enhanced Treg suppressive capacity and lineage-stability. MTHFD2 deficiency also promoted aberrant FoxP3 expression and suppressive activity in Th17 cells. These effects were associated with intracellular accumulation of AICAR, downregulation of the mTORC1 pathway, decreased purine synthesis, and changes in histone methylation. *In vivo* targeting of MTHFD2 protected against both experimental autoimmune encephalomyelitis (EAE) and delayed-type hypersensitivity (DTH). Together these data show that MTHFD2 serves as a metabolic checkpoint in Th17 and Treg cells and highlight the potential of this enzyme as a novel target for anti-inflammatory immunotherapy.

## RESULTS

### Nucleotide synthesis and 1C metabolism is differentially active in CD4 T cell subsets

Rapid cell growth and proliferation following CD4 T cell activation requires production of nucleotides to meet DNA and RNA synthesis demands. We hypothesized that this process is differentially regulated in CD4 subsets. We first analyzed metabolomics data to establish nucleobase levels in CD4 T cells 24 hours post activation with anti-CD3 and anti-CD28 antibodies (Kishton et al., 2016). At this early timepoint, both purine and pyrimidine nucleobase levels were elevated compared to resting cells (**Figure 1A**). At a later timepoint, five days post *in vitro* differentiation, different levels of nucleobases (**Figure 1B**) and purine synthesis intermediates (**Figure 1C**) were measured in Th1, Th17, and Treg subsets (Gerriets et al., 2014). Th17 cells were notable for significantly increased levels of guanine, as well as the purine synthesis intermediate formyl-glycinamide ribonucleotide (FGAR), and the depletion of the downstream intermediate AICAR compared to resting cells.

**Figure 1:**
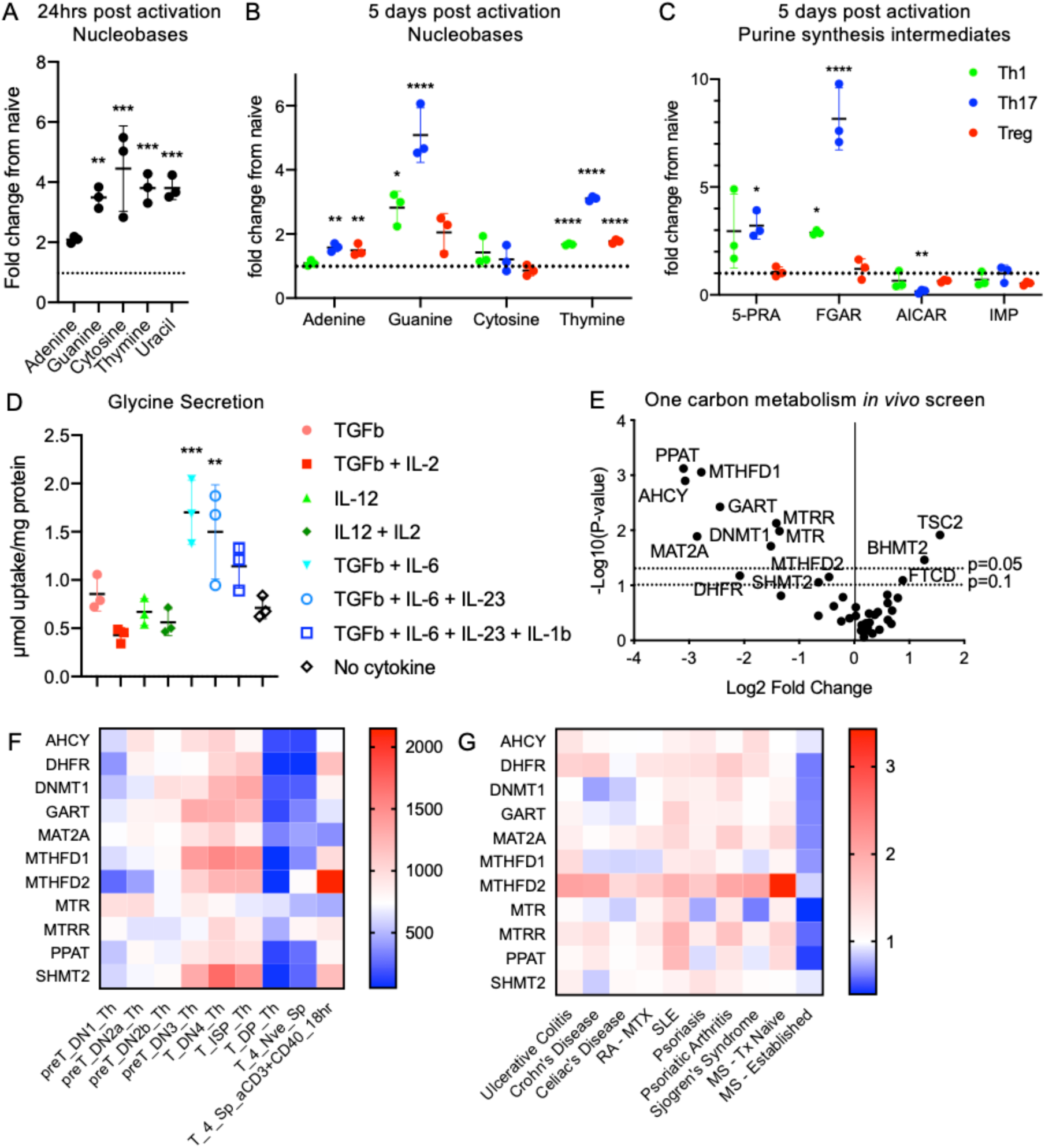
1C metabolism and *de novo* purine synthesis pathways are differentially active in CD4 T cells. (A) Fold change in nucleobase levels in undifferentiated CD4 T cells 24 hours post activation with anti-CD3/anti-CD28 compared to naïve cells measured by MS (one-way ANOVA) (Kishton et al., 2016). (B, C) Fold change in nucleobase (B) and purine synthesis intermediates (C) levels in Th1, Th17, and Treg *in vitro* differentiated cells five days post activation compared to resting cells measured by MS (one-way ANOVA) (Gerriets et al., 2014). (D) Glycine uptake of cells cultured in indicated cytokine conditions for 72 hours post activation measured by <u>^1^H-MRS</u>, corrected for cell proliferation by protein mass (one-way ANOVA). (E) Change in gRNA abundance from *in vivo* CRISPR/Cas9-based screening of one carbon metabolism genes in primary CD4 T cells in a lung-inflammation model (analyzed by MaGECK). (F) mRNA expression of genes identified in (E) during T cell development and activation. Data from ImmGen RNA-seq databrowser. (G) mRNA expression of genes identified in (E) in whole blood of patients with indicated inflammatory disorders (Aune et al., 2017). (RA-MTX = rheumatoid arthritis under methotrexate treatment, SLE = systemic lupus erythematosus, MS-Tx Naïve = multiple sclerosis at time of diagnosis, MS-Established = multiple sclerosis established and under treatment.) See also Figure S1.

*De novo* purine synthesis relies on the folate cycle in 1C metabolism for the two intermediate formylation steps. Metabolism of serine to glycine provides 1C units that are transferred through several reactions in the folate cycle to generate formate. Consistent with upregulation of this pathway and endogenous production of glycine, we found *in vitro* differentiated Th17 cells secrete more glycine than other CD4 subsets (**Figure 1D**) when normalized for total cellular protein (**Supplemental Figure 1A**). This supports the hypothesis that this pathway is particularly active in Th17 cells. Surprisingly, no measurable differences were found in glucose and glutamine uptake or lactate secretion among the CD4 subsets when corrected for proliferation rates by total cellular protein (**Supplemental Figure 1B**).

To further interrogate the dependence of CD4 T cells on enzymes in 1C metabolism and rank-order genes in this pathway for impact on T cell function *in vivo*, we performed an *in vivo* pooled retroviral CRISPR-Cas9 based targeted screen in primary CD4 T cells (**Figure 1E**). CRISPR screening allows systematic evaluation of a list of potential targets that can rank order the effect magnitude of knocking out specific genes. Here, a custom guide RNA (gRNA) library was constructed to target enzymes in 1C metabolism, along with Tuberous Sclerosis Complex 2 (TSC2) as a positive control and non-targeting negative controls. CD4 T cells isolated from ovalbumin-specific OT-II/Cas9 double-transgenic mice were retrovirally transduced with this library and intravenously transferred into Rag1-/- hosts. Mice were then immunized with intranasal ovalbumin to induce lung inflammation. T cells recovered from the lungs of these mice were sequenced and enrichment or depletion were established relative to non-targeting controls. TSC2 gRNA was enriched, supporting the inhibitory role of this protein on T cell proliferation and effector function, with consequent enrichment of TSC2-deleted T cells. In contrast, PPAT, AHCY, MAT2A and MTHFD1 gRNA were significantly depleted and GART, DHFR, DNMT1, MTRR, MTR, SHMT2, and MTHFD2 gRNA were reduced to a lesser extent to indicate these genes play important roles for T cells *in vivo* (**Figure 1E, Supplemental Figure 1C**). MTHFD1 in the cytosol and MTHFD2 in the mitochondrial compartment mediate the synthesis of 10-formylTHF. MTHFD1 loss in particular results in complete ablation of purine biosynthesis, but the function of MTHFD2 can be redundant with MTHFD1. SHMT2 and MTHFD2 lie in tandem in the mitochondrial pathway of the folate cycle. These results indicate *in vivo* dependence of primary CD4 T cells on these genes and 10-formylTHF generation.

We next examined the expression profile of the genes identified from the above *in vivo* CRISPR screen. While coordinately upregulated in developing thymocytes, MTHFD2 was most strongly upregulated in stimulated peripheral T cells (**Figure 1F**). Moreover, in a separate published RNAseq dataset from the whole blood of patients with a variety of inflammatory and autoimmune diseases (Aune et al., 2017), MTHFD2 was consistently overexpressed across multiple conditions including ulcerative colitis, Crohn’s disease, Celiac’s disease, rheumatoid arthritis (RA), systemic lupus erythematosus (SLE), psoriasis, psoriatic arthritis, Sjogren’s syndrome, and MS (**Figure 1G**). In particular, patients with newly diagnosed MS had significantly elevated MTHFD2 expression in their whole blood compared to healthy donors as well as MS patients undergoing therapy with disease remission (**Supplemental Figure 1D**).

### MTHFD2 is highly upregulated in CNS-infiltrating CD4 T cells in the setting of EAE and in activated CD4 T cells in vitro

To measure the expression of MTHFD2 in inflammatory lesions *in vivo*, CD4 T cells from myelin-specific T-cell receptor (TCR) transgenic 2D2 mice were activated and adoptively transferred into Rag1-/- mice to induce EAE. After mice began exhibiting hind-leg paralysis, the spinal cords were analyzed by immunohistochemistry. The cauda equina segment of the spinal cord of mice that received 2D2 T cells showed enrichment of MTHFD2 and CD3 positive cells that was absent in the control mice (**Figure 2A**). To directly determine if CD4 T cells upregulated MTHFD2 in inflamed lesions, cells from spleen and spinal cord of mice subjected to a myelin oligodendrocyte glycoprotein (MOG) and pertussis toxin (PTX) induced model of EAE were collected for flow cytometry. CD4 T cells infiltrating the spinal cord of these mice showed overexpression of MTHFD2 compared to matched splenic CD4 T cells from control and EAE mice as measured by flow cytometry (**Figure 2B**) and by RT-qPCR levels of purified CD4 T cells (**Figure 2C**).

**Figure 2:**
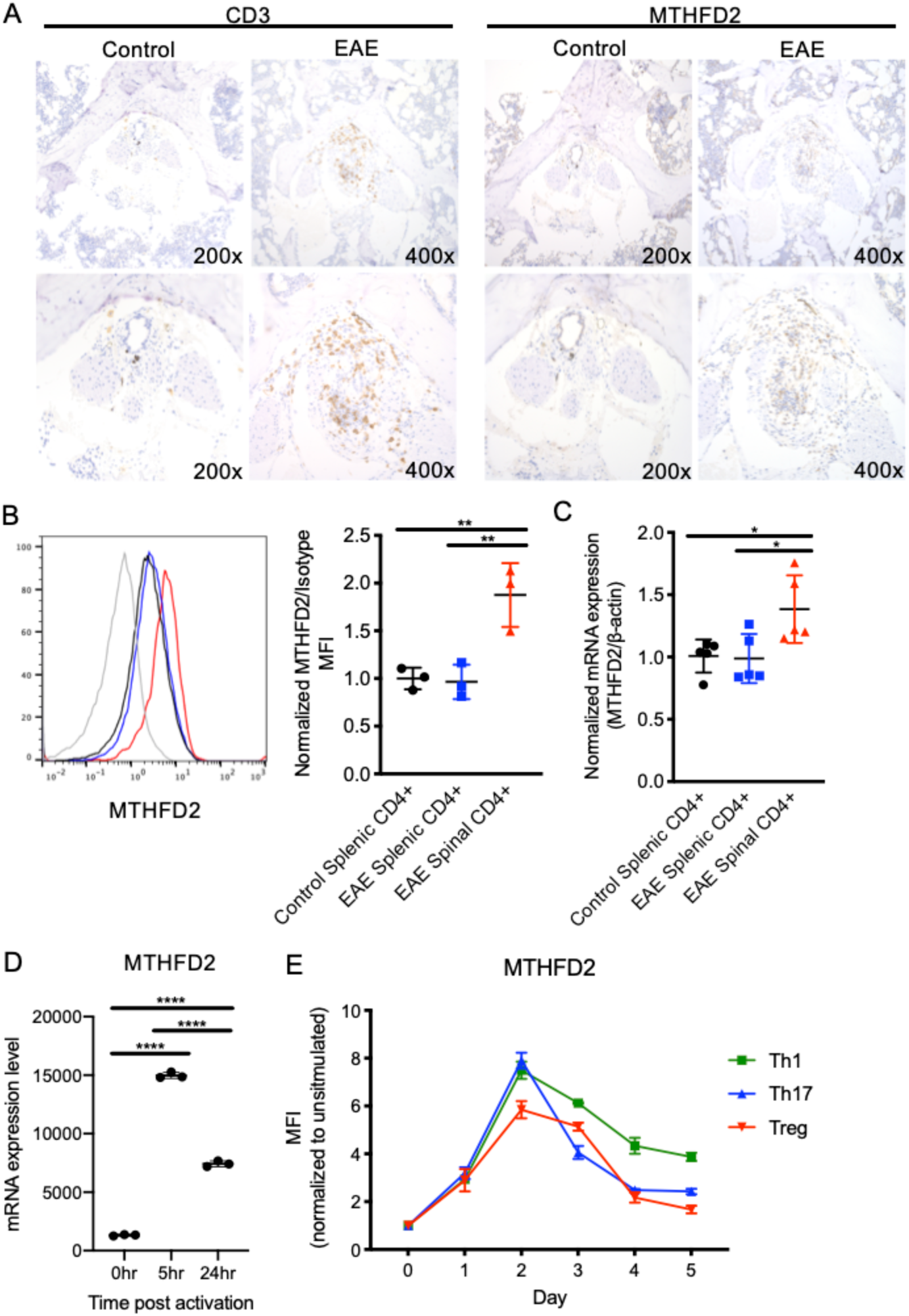
MTHFD2 is upregulated in CNS-infiltrating T cells in Experimental Autoimmune Encephalomyelitis (EAE) *in* vivo and in pathogenic Th17 cells *in vitro*. (A) Immunohistochemistry showing CD3 and MTHFD2 staining in the cauda equina of mice with symptomatic EAE. (B-C) Relative mean fluorescence intensity (MFI) normalized to isotype (B) and relative mRNA expression normalized to β-actin (C) of MTHFD2 in CD4 T cells isolated from spleen and spinal cord of mice with symptomatic EAE and spleen of control mice, (one-way ANOVA). (D) MTHFD2 mRNA expression in undifferentiated CD4 T cells at 0, 5, and 24 hours post activation with anti-CD3/anti-CD28 (one-way ANOVA). (E) Relative MTHFD2 MFI over five days post activation in Th1, Th17, and Treg cells, normalized to expression in resting cells. See also Figure S1.

To characterize the kinetics of MTHFD2 expression in CD4 cells, change in mRNA expression in response to *in vitro* activation was measured. Activation induced robust upregulation of MTHFD2 mRNA levels by 5 hours, and expression began to fall by 24 hours (**Figure 2D**). *In vitro* differentiated Th1 and Th17 cells that drive EAE pathogenesis were found to attain the highest level of MTHFD2 protein expression 48 hours post activation (**Figure 2E**). Thus, MTHFD2 is rapidly upregulated with activation in CD4 T cells and is associated with pathogenic CD4 T cell activity in the setting of EAE.

### CD4 T cell subsets differentially require MTHFD2

To test the role of MTHFD2 in CD4 T cell subset activation, differentiation, proliferation, and function, CD4 cells were activated *in vitro* in the presence of cytokines for optimal Th1, Th17, and Treg differentiation together with vehicle or an MTHFD2 inhibitor (MTHFD2i; DS18561882) (Kawai et al., 2019). After 72 hours of incubation with the inhibitor, all subsets were found to have significantly reduced cell numbers, although only Th17 cells had statistically significantly reduced viability (**Figure 3A**). Activation, as measured by CD25 expression, was also reduced in all subsets (**Figure 3B**). Lineage-characterizing transcription factor expression was reduced in Th1 cells (T-bet), but unchanged in Th17 cells (RORγt) and Treg cells (FoxP3) (**Figure 3C**). To measure effector T cell function, cells were stimulated with PMA and ionomycin. In the presence of the MTHFD2i, significantly fewer Th1 cells expressed IFNγ, and fewer Th17 cells expressed IL-17 (**Figure 3D**). These data indicate MTHFD2 function is required for maximal CD4 T cell proliferation and cytokine production.

**Figure 3:**
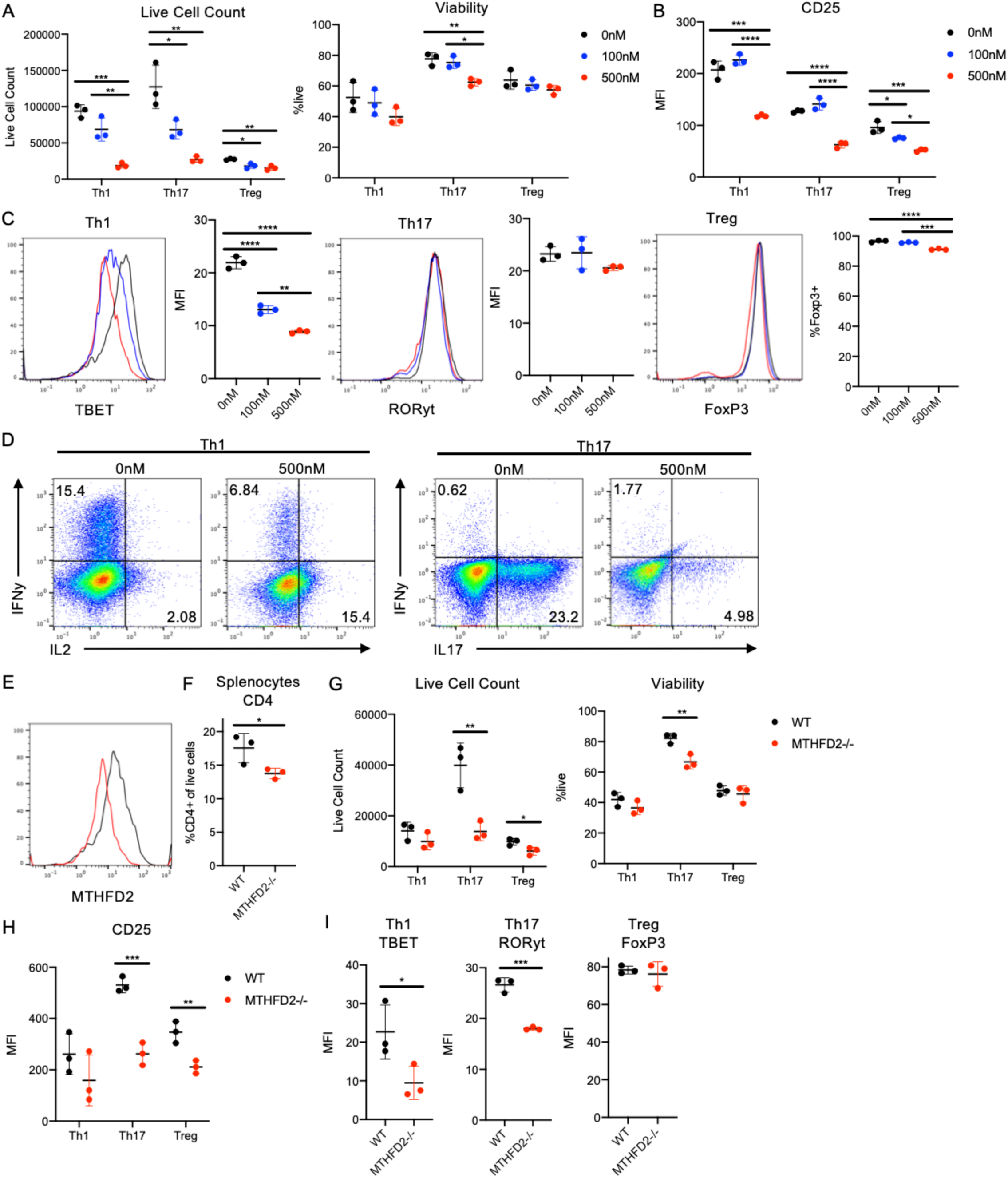
MTHFD2 deficiency impairs CD4 T cell proliferation and function. (A-C) Live cell count (left) and viability (right) (A), CD25 expression (B), and transcription factor expression (T-bet in Th1, left; RORγt in Th17, middle; FoxP3 in Treg, right) (C) in Th1, Th17, and Treg cells treated with 0, 100, or 500nM DS18561882 for 72 hours post activation with anti-CD3/anti-CD28 (one-way ANOVA). (D) Cytokine expression in Th1 (IFNγ and IL2, left) and Th17 (IFNγ and IL17, right) cells treated with 0 or 500nM DS18561882 for four days post activation and stimulated with PMA and ionomycin for four hours. Representative of 3 biological replicates. (E) MTHFD2 expression in activated CD4 T cells isolated from MTHFD2^fl/fl^ CD4-Cre^+^ (MTHFD2^-/-^) and MTHFD2^fl/fl^ CD4-Cre^-^ (WT) control littermates. (F) Frequency of CD4 T cells in the spleen of treatment-naïve MTHFD2^-/-^ and WT littermates (unpaired t-test). (G-I) Live cell count (left) and viability (right) (G), CD25 expression (H), and transcription factor expression (T-bet in Th1, left; RORγt in Th17, middle; FoxP3 in Treg, right) (I) four days post activation in Th1, Th17, and Treg cells from MTHFD2^-/-^ and WT littermates (unpaired t-test).

To determine whether these pharmacological effects were recapitulated genetically, we generated a MTHFD2^fl/fl^ mouse strain and crossed it with CD4-Cre transgenic mice for several generations to achieve conditional knockout. As anticipated, Cre^+^ (MTHFD2^-/-^) CD4 T cells expressed lower levels of MTHFD2 compared to cells from Cre^-^ (WT) littermates (**Figure 3E**). At baseline, MTHFD2^fl/fl^ CD4-Cre^+^ mice had slightly fewer CD4 T cells in the spleen (**Figure 3F**). Upon *in vitro* activation and differentiation, cell numbers and viability were significantly lower in MTHFD2^-/-^ Th17 cells (**Figure 3G**). Proliferation was also slightly dampened in MTHFD2^-/-^ Treg cells. In addition, CD25 expression levels were decreased in Th17 and Treg cells, and trended lower in Th1 cells (**Figure 3H**). Finally, MTHFD2^-/-^ Th1 and Th17 cells had lower expression of T-bet and RORγt, respectively. FoxP3 in Treg cells, however, was unchanged (**Figure 3I**). Differences in results between the pharmacological and genetic approaches may be attributed to the timing and completeness of enzyme function loss during development and potential compensatory mechanisms. These experiments collectively show a requirement for MTHFD2 in CD4 T cell activity.

### MTHFD2i induces FoxP3 expression in pathogenic Th17 cells and increases Treg lineage generation and stability

To further interrogate the inhibitory effects of MTHFD2 deficiency on Th1 and Th17 cells, FoxP3 was measured in each condition. Interestingly, treatment with MTHFD2i induced aberrant upregulation of FoxP3 in an inhibitor dose-dependent manner (**Figure 4A**). While most prominent in Th17 cells, FoxP3 was also upregulated in Th1 cells to a modest extent. A similar trend was observed in the MTHFD2^fl/fl^ CD4-Cre^+^ mice, but with FoxP3 only induced in Th17 cells (**Figure 4B**). Consistent with a functional response to aberrant expression of FoxP3, MTHFD2i-treated Th17 cells also exhibited suppressive capacity when co-cultured with CD8 T cells (**Figure 4C**). These data suggest that MTHFD2i promotes FoxP3 and Treg-like phenotypes in Th17 cells.

**Figure 4:**
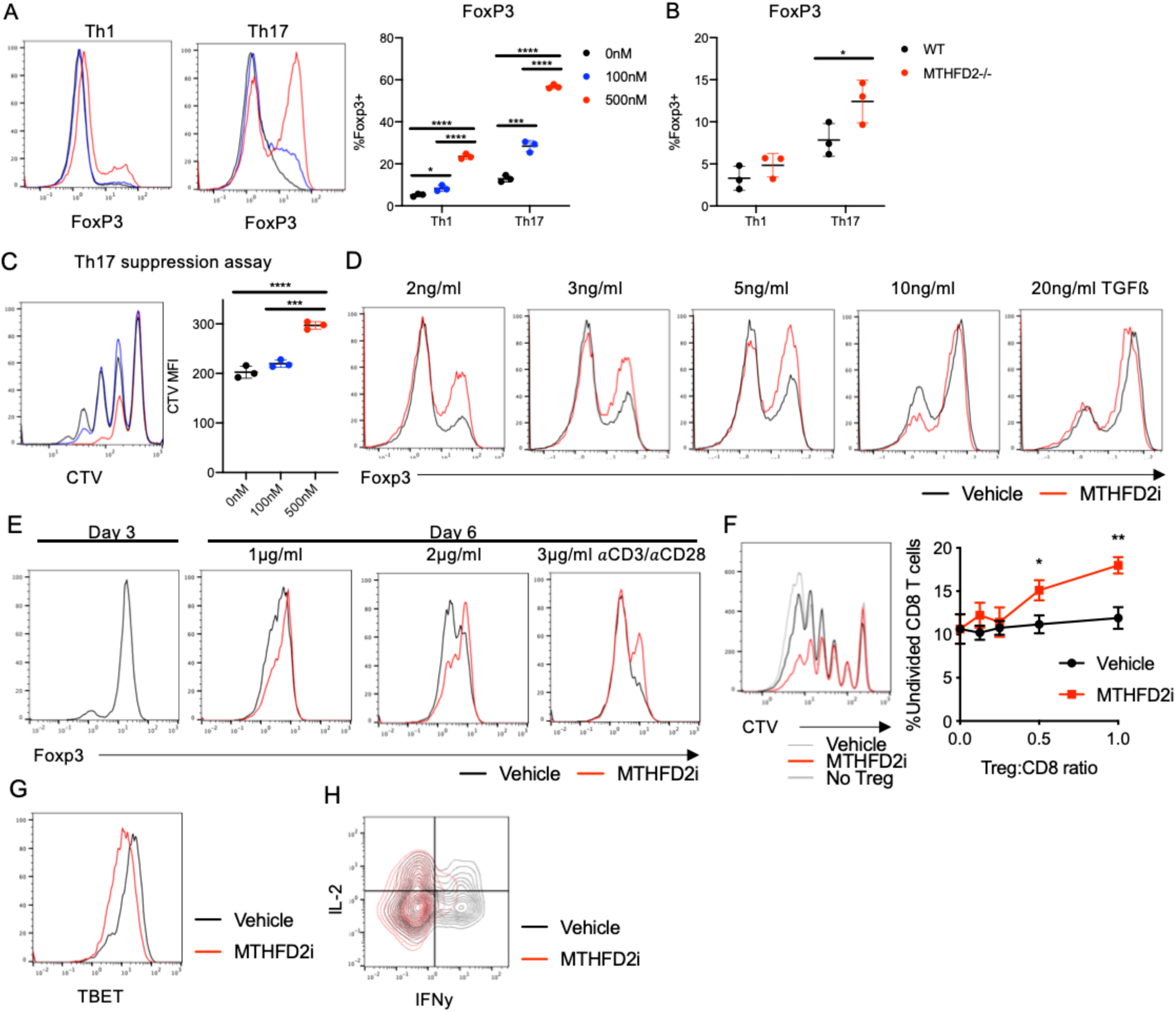
MTHFD2 deficiency stabilizes FoxP3 expression in Th17 and Treg cells. (A) FoxP3 expression in Th1 and Th17 cells 72 hours post activation with anti-CD3/anti-CD28 treated with 0, 100, or 500nM DS18561882 (one-way ANOVA). (B) FoxP3 expression in Th1 and Th17 cells 4 days after activation from MTHFD2^-/-^ and WT littermates (unpaired t-test). (C) Th17 suppression assay measuring aberrant suppressive capacity of Th17 cells treated with 0, 100, or 500nM DS18561882. Pre-treated Th17 cells were co-cultured with CD8 T cells stained with CTV to measure proliferation upon activation with anti-CD3/anti-CD28 (one-way ANOVA). (D) FoxP3 expression in induced Treg cells differentiated with low potency TGFβ at a range of concentrations treated with either vehicle or 500nM DS18561882. (E) FoxP3 expression in induced Treg cells cultured for three days post activation (left). On day 3, cells were restimulated with varying concentrations of anti-CD3/anti-CD28 with IL-2 and either vehicle or 2.5μM Raze 1459. Change in FoxP3 expression by day 6 (right). (F) Treg suppression assay showing proliferation of CTV-stained CD8 T cells co-cultured with Treg cells from day 6 of (E) at different ratios to measure residual suppressive capacity. Treg cells were pre-treated with either vehicle or 2.5μM Raze 1459 and restimulated on 3μg/ml anti-CD3 and 5μg/ml anti-CD28 to induce FoxP3 loss prior to suppression assay. (G-H) Th1-phenotypic T-bet (G) and IFNγ (H) expression in same cells from (E) on day 6. Vehicle or 2.5μM Raze 1459.

MTHFD2 inhibition also directly enhanced differentiation and stability of induced Treg cells. In contrast to the optimized conditions above, fewer cells expressed FoxP3 when Treg cells were cultured with lower levels of TGFβ (**Figure 4D**). Treg cells were thus treated with MTHFD2i in the presence of a range of TGFβ concentrations. At every concentration except at the highest tested, FoxP3 expression increased with MTHFD2i treatment **(Figure 4D).** Induced Treg cells can have a low degree of lineage stability, and stimulated Treg cells can lose FoxP3 expression and instead acquire effector T cell attributes. However, when induced Treg cells were restimulated in the presence of MTHFD2i, they were better able to maintain FoxP3 expression over a range of anti-CD3 concentrations (**Figure 4E**). Consistent with elevated FoxP3 expression, MTHFD2i-treated Treg cells also retained greater suppressive capacity (**Figure 4F).** At the same time, MTHFD2i prevented the upregulation of T-bet and IFNγ seen in the restimulated vehicle-treated Treg cells (**Figure 4G and H**). Therefore, MTHFD2i enhanced Treg generation and reduced plasticity to other subsets in restimulated Treg cells.

### MTHFD2 deficiency in human CD4 T cells decreases Th17 proliferation and improves Treg lineage stability

To test the translatability of the above findings in human cells, CD4 T cells isolated from human PBMCs were *in vitro* differentiated into Th1, Th17, and Treg cells and treated with either vehicle or MTHFD2i from the time of activation. In all subsets, proliferation was decreased during activation (**Figure 5A**). Of the subsets, Th17 showed the greatest sensitivity when treated with MTHFD2i post activation (**Figure 5B**). Next, FoxP3 expression was measured in Th17 and Treg cells. In both subsets, FoxP3 expression was higher in the MTHFD2i treated condition than the vehicle (**Figure 5C**). In an experiment parallel to **Figure 4D**, human Treg cells were tested for FoxP3 lineage stability in the presence or absence of the inhibitor. Consistent with the murine data, MTHFD2i treated cells expressed higher levels of FoxP3 and Helios, a marker of Treg cells with high suppressive capacity and stable Foxp3 expression (Fuhrman et al., 2015; Thornton and Shevach, 2019), compared to vehicle 48hrs post re-stimulation (**Figure 5D**). Finally, the above findings were replicated genetically using shRNA to silence MTHFD2. Cells transfected with MTHFD2 shRNA were confirmed to have knockdown of MTHFD2 protein expression compared to those transduced with control shRNA (**Figure 5E).** All cell subsets proliferated less with MTHFD2 shRNA, with the greatest reduction in Th17 cells as determined by Ki67 staining (**Figure 5F**). Thus, both the loss of proliferative capacity and stabilization of FoxP3 expression in Th17 and Treg cells can be recapitulated in primary human CD4 T cells.

**Figure 5:**
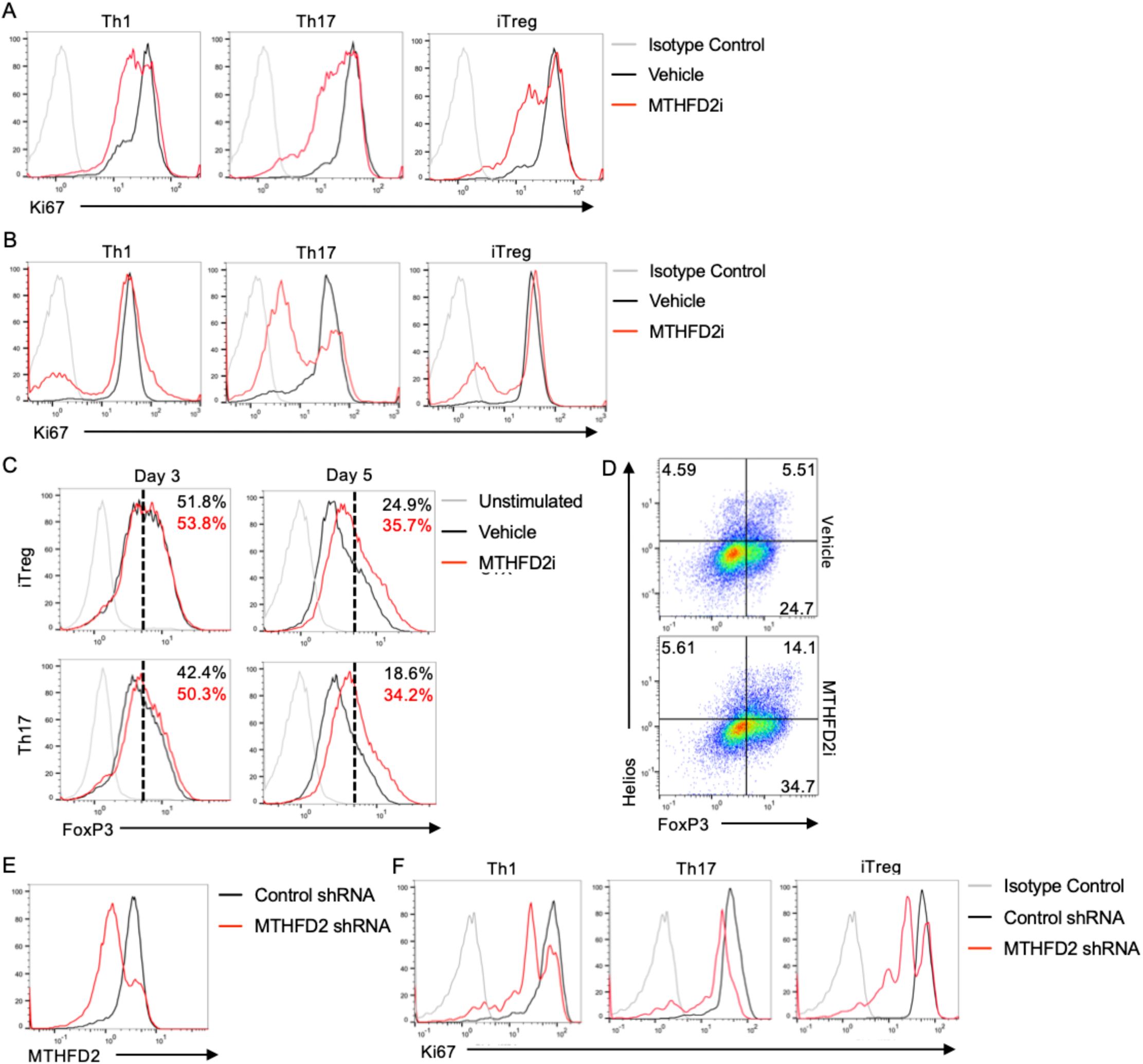
MTHFD2 deficiency impairs proliferation and stabilizes FoxP3 expression in human primary CD4 T cells. (A-B) Ki67 expression in human Th1, Th17, and iTreg cells treated with either vehicle or 2μM DS18561882 from day 0 to 3 (A) or from day 3 to 5 (B) post activation with anti-CD3/anti-CD28. (C) FoxP3 expression in Th17 and iTreg cells treated with either vehicle or 2μM DS18561882 on days 3 and 5 post activation. (D) FoxP3 and Helios expression in iTreg cells on day 5, treated with either vehicle or 2μM DS18561882 from day 3. (E) MTHFD2 protein expression in CD4 T cells transduced with either control or MTHFD2 shRNA. (F) Ki67 expression in Th1, Th17, and Treg cells treated with control or MTHFD2 shRNA. All plots representative of three experiments.

### MTHFD2 activity regulates mTORC1 activity in Th1 and Treg cells, and methylation levels in Th17 cells

As MTHFD2 is required for the maintenance of the mitochondrial formate pool, we measured formate exchange in response to MTHFD2i (Ma et al., 2017). As expected, cells treated with MTHFD2i significantly decreased formate secretion into the media (**Figure 6A**). The 1C unit used for formate synthesis is largely derived from the conversion of serine to glycine. With MTHFD2i, intracellular serine accumulated, and cells switched from glycine export to import, further supporting an on-target effect of the MTHFD2i (**Figure 6B and C**). We next tested whether the provision of formate in the media to rescue the enzymatic function of MTHFD2 could reverse the effects of the inhibitor. Indeed, 1mM formate was sufficient to restore proliferation as well as activation in MTHFD2i treated cells (**Figure 6D**). Notably, addition of 1mM formate reduced FoxP3 levels to baseline in Treg cells differentiated with low levels of TGFβ (**Figure 6E**).

**Figure 6:**
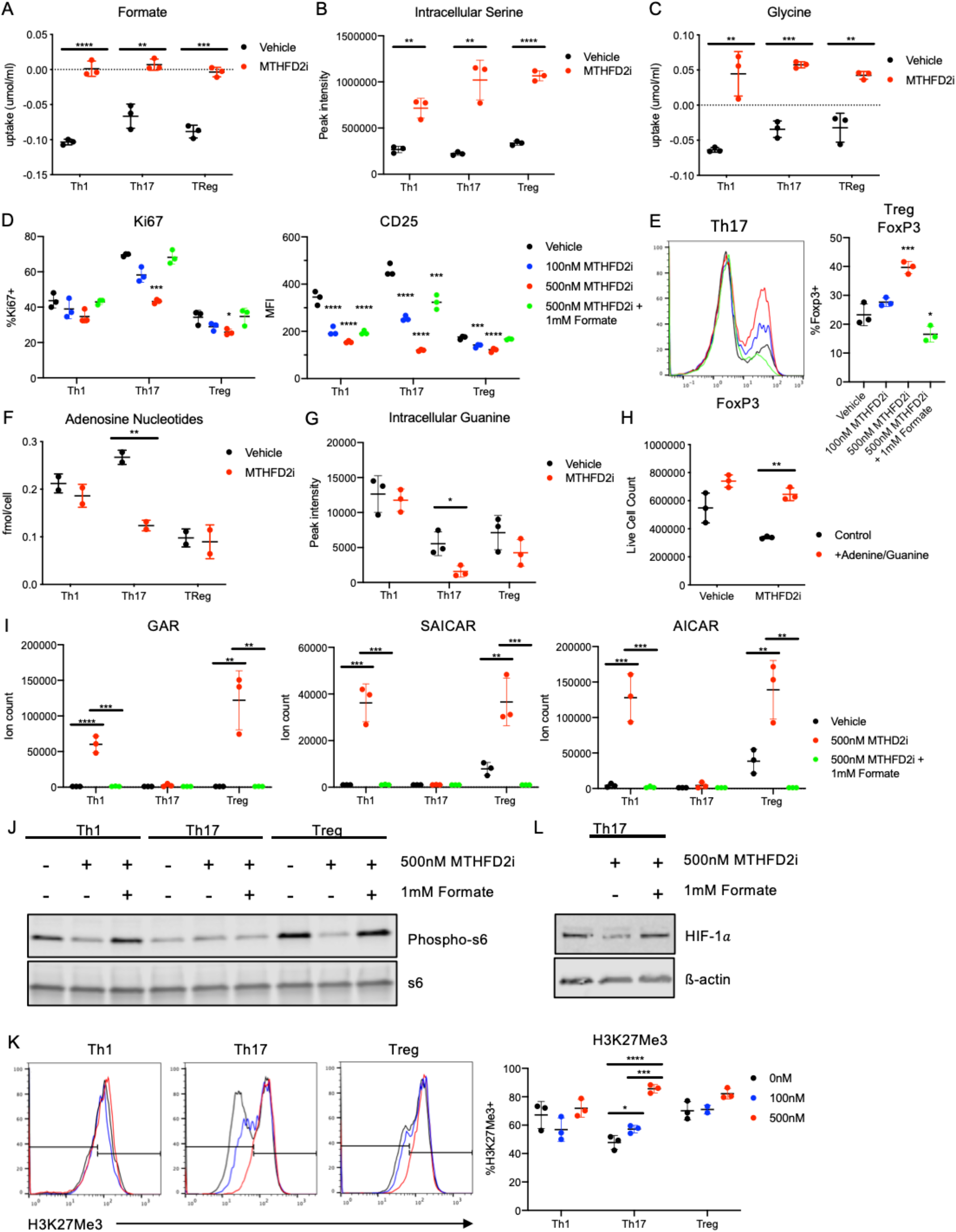
MTHFD2i effects are mediated by formate depletion resulting in accumulation of purine synthesis intermediates, decreased mTORC1 activity, and altered histone methylation levels. (A-C) Formate uptake measured by <u>^1^H-MRS</u> (A), intracellular serine levels measured by MS (B), and glycine uptake measured by <u>^1^H-MRS</u> (C) in Th1, Th17, and Treg cells activated and differentiated for 72 hours and then treated with either vehicle or 2.5μM Raze 1459 for 48hrs (unpaired t-test). (D-E) Ki67 (left) and CD25 (right) MFI in Th1, Th17, Treg cells (D) and FoxP3 expression in Treg cells differentiated in low potency TGFβ (E) treated with 0nM, 100nM, 500nM DS18561882, or 500nM DS18561882 with 1mM formate for 72 hours post activation (one-way ANOVA). (F-G) Intracellular adenosine nucleotide measured by ^1^H-MRS (A) and guanine measured by MS (B) levels in Th1, Th17, and Treg cells treated with either vehicle or 2.5μM Raze 1459 for 48hrs (unpaired t-test). (H) Th17 cell numbers after treatment with either vehicle or 2.5μM Raze 1459 in base media or media supplemented with adenine and guanine (unpaired t-test). (I-J) MS measurement of GAR, SAICAR, and AICAR (I) and immunoblot of phospho-S6 (J) in Th1, Th17, and Treg cells treated with 0nM, 100nM, 500nM DS18561882, or 500nM DS18561882 with 1mM formate (one-way ANOVA). (K) H3K27-trimethylation levels in Th1, Th17, and Treg cells treated with 0nM, 100nM, or 500nM DS18561882 measured by flow cytometry (one-way ANOVA). (L) Immunoblot of HIF-1α in Th17 cells treated with vehicle, 500nM DS18561882, or 500nM DS18561882 with 1mM formate. See also Figure S2.

Consequent changes in *de novo* purine synthesis were investigated by measuring levels of intracellular adenosine nucleotides and guanine. With extended MTHFD2i treatment of 48 hours, these purine products were only depleted in Th17 cells and not Th1 or Treg cells (**Figure 6F and G)**. This finding was reproduced with a second MTHFD2i (Kawai et al., 2019) (**Supplemental Figure 2A**). These changes were sufficient to reduce proliferation, as media supplemented with adenine and guanine was sufficient to rescue cell numbers in MTHFD2i-treated Th17 cells (**Figure 6H).** To determine immediate effects of MTHFD2i on purine synthesis pathway intermediates, cells were cultured for 72 hours before transient exposure to MTHFD2i for four hours. This short timeframe was chosen to avoid any secondary effects including compensatory changes. In immortalized cancer cells, MTHFD2 deficiency leads to accumulation of glycineamide ribonucleotide (GAR), 1-(phosphoribosyl) imidazolecarboxamide (SAICAR), and AICAR, which are upstream of 10-formylTHF-mediated formylation steps (Ducker et al., 2016). Here, transient exposure resulted in significant accumulation of GAR, SAICAR, and AICAR in Th1 and Treg cells but not Th17 cells (**Figure 6I**). This effect was fully rescued by addition of 1mM formate, supporting an on-target enzymatic effect of MTHFD2i. AICAR is an established regulator of the mTORC1/AMPK axis (Kim et al., 2016), a major driver of metabolic reprogramming. In accordance with the mass spectrometry results, phosphorylation of the mTORC1 target ribosomal protein S6 (phospho-S6) was decreased with MTHFD2i in Th1 and Treg cells but not Th17 cells. Again, phospho-S6 levels were restored by addition of formate (**Figure 6J**).

It remained unclear how MTHFD2-deficiency impaired Th17 cells. The 1C metabolism pathway and MTHFD2 activity can contribute to cellular redox balance through conversion of NAD+ to NADH and NADP+ to NADPH (Shin et al., 2017). It also feeds into the transsulfuration pathway and subsequent glutathione synthesis. Both total cellular reactive oxygen species (ROS) and mitochondrial superoxide levels, however, were unchanged with MTHFD2i treatment (**Supplemental Figure 2B**). Next, possible changes in global methylation levels as a consequence of MTHFD2i were investigated. Folate and methionine cycles are required for production of S-adenosyl methionine (SAM), which serves as the methyl donor in DNA and protein methylation reactions. Histone methyltransferases (HMTs) convert SAM to S-adenosylhomocysteine (SAH). HMT function depends on methionine and SAM concentrations and is inhibited by SAH accumulation. Thus, the ratio SAM/SAH is thought to be indicative of general cellular methylation potential (Mentch et al., 2015). Interestingly, *in vitro* differentiated Th1, Th17, and Treg cells were found to have significantly different levels of SAM and SAH (Gerriets et al., 2014). Th17 cells in particular had high levels of SAH, while Treg cells had high levels of SAM. Consequently SAM/SAH ratio was dramatically lower in Th17 cells compared to Treg cells (**Supplemental Figure 2C**). In response to MTHFD2i, H3K4- and H3K27-trimethylation levels were both specifically increased in Th17 cells but not Th1 and Treg cells in a dose-dependent manner (**Figure 6K, Supplementary Figure 2D**). Associated with this increase in methylation, the expression of HIF-1α, a key transcription factor required for Th17 differentiation and attenuation of Treg differentiation (Shi et al., 2011), was decreased with MTHFD2i treatment and restored with addition of formate (**Figure 6L**). Specifically, HIF-1α has been shown to activate transcription of Th17 genes and flag FoxP3 for protein degradation (Dang et al., 2011). In contrast, the expression of HIF-2α, which supports Treg cell function (Hsu et al., 2020), is slightly elevated with treatment (**Supplementary Figure 2E**). The contrasting expression changes for HIF-1α and HIF-2α suggest that this response was not due to hypoxia and support the FoxP3 upregulation phenotype observed in Th17 cells in response to MTHFD2i.

### MTHFD2i in vivo reduces severity of Experimental Autoimmune Encephalomyelitis (EAE) and Delayed-Type Hypersensitivity (DTH)

The efficacy of MTHFD2 as a therapeutic target was next tested *in vivo*. First, EAE was induced by immunization with MOG_35-55_ peptide and CFA emulsion on day 0, and PTX on day 0 and 2. From day 4 onward, mice were treated daily with intraperitoneal injection of either DMSO or the orally available dual MTHFD1/2 inhibitor LY345899 (Gustafsson et al., 2017). Inhibitor treatment led to significantly reduced disease severity and cumulative clinical score compared to vehicle (**Figure 7A and B**). Mice were sacrificed on day 26 and spinal cord infiltrating cells were collected for immune profiling. Notably, significantly fewer CD45+ cells infiltrated into the spinal cord of MTHFD2i treated mice (**Figure 7C**) along with fewer CD4+ and CD8+ T cell, although the frequencies of these populations were unchanged (**Figure 7D and E**). MTHFD2i treated mice also had decreased CNS infiltration of RORγt^+^ CD4 cells (**Figure 7F)**, decreased frequency of IFNγ^+^IL17^+^ doublepositive cells, and increased frequency of IL17^+^ single-positive cells (**Figure 7G**), suggesting a possible switch to a less pathogenic phenotype. Finally, FoxP3^+^CD25^+^ CD4 cell numbers were reduced and IFNγ^+^ CD4 cell numbers showed a trend towards lower level in treatment conditions (**Supplemental Figures 3A and B**).

**Figure 7:**
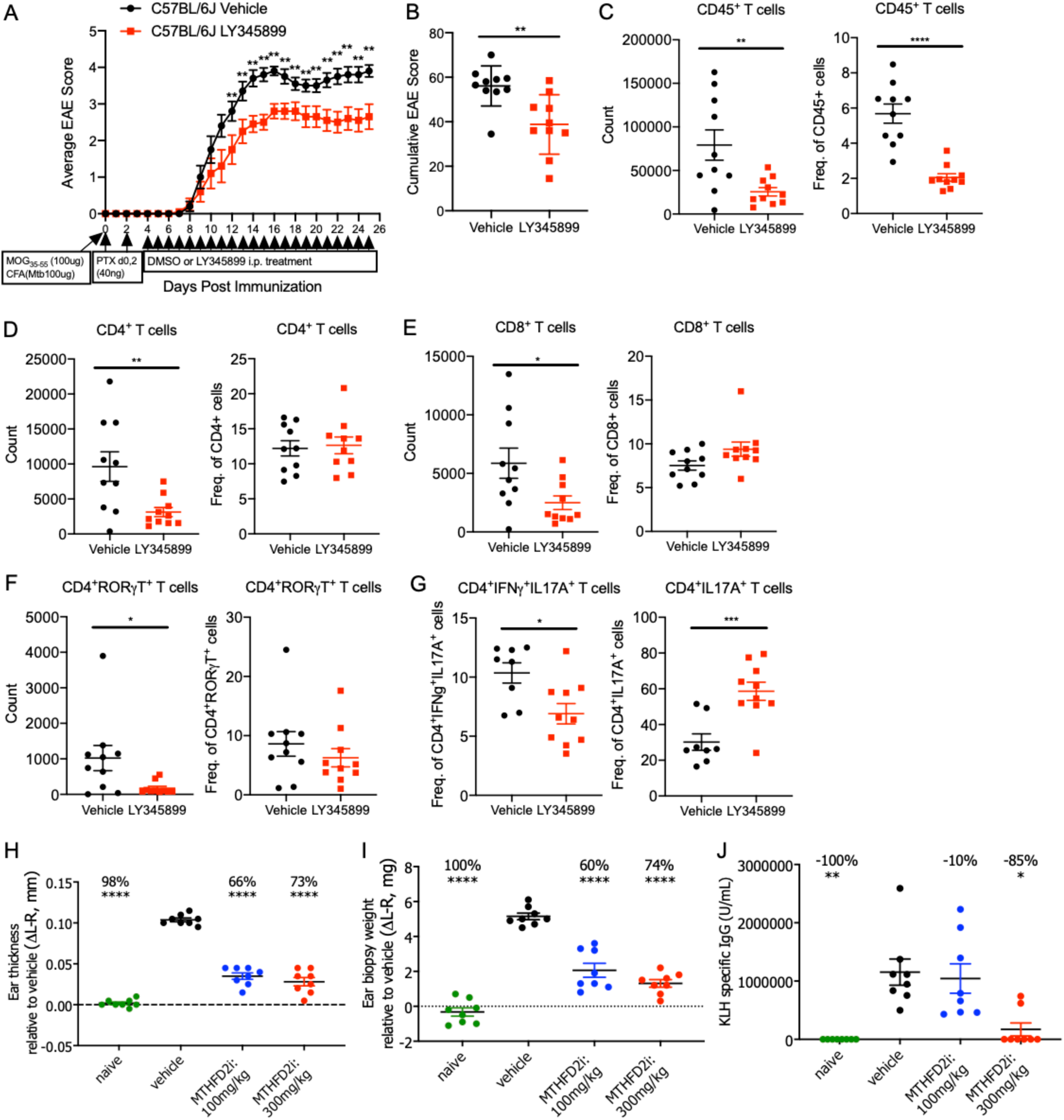
*In vivo* MTHFD2i reduces disease severity in EAE and Delayed-Type Hypersensitivity (DTH) models. (A-B) Average clinical score overtime (n=10, mean±SEM, multiple t-tests) (A) and cumulative score (Mann-Whitney test) (B) in mice immunized with MOG_35-55_/CFA and PTX and injected intraperitoneally (i.p.) daily with vehicle or LY345899 at 10mg/kg. (C-F) Fraction and total number of CD45+ (C), CD4+ (D), CD8+ (E), and RORγt^+^ T cells in the spinal cord of mice from (A) on day 26 post immunization (unpaired t-test). (G) Fraction of IFNγ^+^IL17A^+^ double-positive and IL17A^+^ single-positive CD4 T cells in the spinal cord (unpaired t-test). See also Figure S3. (H-I) Relative ear thickness (H) and ear biopsy weight (I) 10 days post immunization with KLH/CFA and 3 days post challenge with KLH (one-way ANOVA). Mice were treated twice daily with PO vehicle, 100, or 300mg/kg DS18561882 (mean±SEM). (J) KLH-specific IgG levels on day 10 of KLH-induced DTH (mean±SEM, Kruskal-Wallis test). See also Figure S4.

The role of MTHFD2 was also tested in a T cell dependent delayed type hypersensitivity model using the selective MTHFD2 inhibitor DS18561882. Mice were immunized and challenged on the ear with Keyhole limpet hemocyanin (KLH) to induce local inflammation and were treated with vehicle or DS18561882. Treatment showed no overt toxicity, as treated animals maintained their weight (**Supplemental Figure 4A**). Importantly, ear thickness and weight were each increased in control DTH mice but not in inhibitor-treated animals, indicating a protection from inflammation upon MTHFD2 inhibition (**Figure 7H and I, Supplemental Figure 4B**). KLH-specific IgG was also reduced at the highest dose of inhibitor (**Figure 7J**). Together these data show that MTHFD2 can be targeted *in vivo* to reduce inflammation and prevent immune-related disease.

## DISCUSSION

T cell activation, differentiation, and function require appropriate metabolic reprogramming to meet the cells increased demands for energy, biosynthetic building blocks, and signaling molecules. In this study, we investigated the function of nucleotide and 1C metabolism in primary CD4 T cells. Using a combination of *in vivo* primary T cell CRISPR-based screening and gene expression data, MTHFD2 was identified as a potential target for antiinflammatory therapies. MTHFD2 inhibition or deficiency generally led to decreased CD4 T cell proliferation. Notably, this was associated with induction of FoxP3 expression and suppressive function in Th17 cells, as well as increased generation and lineage stabilization of Treg cells. These data suggest that MTHFD2 may function as a metabolic checkpoint in the Th17/Treg axis, with MTHFD2i skewing the balance from a pathogenic to a more anti-inflammatory phenotype. Indeed, MTHFD2i ameliorated disease severity in two separate *in vivo* models of autoimmunity and hypersensitivity. These results provide evidence of MTHFD2 as an efficacious disease-modifying target in settings of T-cell driven inflammatory diseases.

1C metabolism consists of serine-glycine metabolism, the folate cycle, and the methionine cycle and is central to several processes including *de novo* purine synthesis, methyl-donor generation, and redox regulation (Yang and Vousden, 2016). 1C metabolism is robustly engaged with TCR stimulation (Tan et al., 2017) and is required to support T cell expansion (Ma et al., 2017). The folate cycle is compartmentalized into cytosolic and mitochondrial components. In the cytosol, MTHFD1 interconverts 5,10-methyleneTHF, 10-formylTHF, and formate. In the mitochondria, the same reactions are accomplished by MTHFD2 and MTHFD1L. These resulting intermediates are used to support formylation steps in *de novo* purine synthesis. MTHFD1 and MTHFD2 can also modulate redox state through NAD(H) and NADP(H) generation (Ducker and Rabinowitz, 2017). While some cell types display flexibility in switching to the cytosolic source under settings of mitochondrial pathway dysfunction (Ducker et al., 2016), T cells have been previously proposed to predominantly depend on the mitochondrial pathway to provide 1C units and reductive species (Ron-Harel et al., 2016). T cells have also been shown to rely on serine uptake and *in vivo* immune responses can be modulated by dietary serine levels. The effects of serine starvation are mediated by limiting purine biosynthesis even in the presence of intact salvage pathways and can be bypassed by provision of glycine and formate (Ma et al., 2017). Our data add to these findings and point to MTHFD2 as a critical regulator that influences both CD4 T cell proliferation and differentiation.

While 1C metabolism plays a broad role in cell metabolism, we found some functions were distinct to select T cell subsets. All human and mouse T cells proliferated to a lesser extent with MTHFD2 deficiency, indicating a shared function for MTHFD2 to support T cell growth and division. The effects of MTHFD2 deficiency on T cell differentiation and effector function, however, differed in each subset tested. Th1 cells showed impaired differentiation with reduced induction of T-bet and decreased cytokine production. Th17 cells, which may have the greatest flux through 1C metabolism pathways based on glycine secretion, also had altered differentiation and decreased cytokine production. While RORγt was not altered, MTHFD2-deficient Th17 cells upregulated the Treg transcription factor, FoxP3, and gained an ability to suppress proliferation of activated CD8 T cells. In contrast, Treg cells exhibited increased differentiation and lineage stability. It is now well established that each of these subsets can have unique metabolic requirements (Bantug et al., 2018; Buck et al., 2015). Our data show that MTHFD2 is also selectively required for Teff while promoting Treg function and fates. It is notable that these outcomes differ from GLUT1, GLS, or ASCT2 deficiencies that impair or alter Teff differentiation while having modest effects to directly promote Treg differentiation (Johnson et al., 2018; Macintyre et al., 2014; Nakaya et al., 2014). The trans-differentiation of Th17 to Treg cells and the increased generation and lineage stability of Treg cells was also most evident with MTHFD2 inhibition.

These distinct T cell fates appear dependent on the metabolic activity of MTHFD2 as the effects of MTHFD2 deficiency could be metabolically rescued with formate or purine nucleobases. While we found no significant effect of altered redox balance with MTHFD2 deficiency, altered *de novo* purine synthesis appears critical. MTHFD2i led to decreased purine levels, particularly in Th17 cells, while also inducing accumulation of the purine synthesis intermediates GAR, SAICAR, and AICAR in Th1 and Treg cells. These intermediates require formylation for further biosynthesis and the effects were rescued when the MTHFD2-derived metabolite formate was added to the media, supporting an on target biochemical mechanism for MTHFD2 inhibition. While differential accumulation of intermediates and depletion of purines were observed in different CD4 subsets, this is likely due to differences in the timing of activation, MTHFD2 activity, and compensatory mechanisms through MTHFD1 or purine salvage pathways. *De novo* nucleotide synthesis to support DNA and RNA synthesis is essential for T cell proliferation (Quéméneur et al., 2003, 2004). Thus, a component of the mechanism by which MTHFD2 inhibits T cell expansion appears to be through insufficient generation of nucleotides.

AICAR accumulation is also notable as this metabolite can have multiple regulatory functions. In cancer cells, AICAR can have cytotoxic effects and increase sensitivity to chemotherapeutics and radiotherapy. These effects are often mediated by AMPK as AICAR is an adenosine analog and an established AMPK activator (Rae and Mairs, 2019; Su et al., 2019). Depending on the setting, however, AICAR may have AMPK-independent effects (Dembitz et al., 2019). Moreover, mTORC1 itself is acutely sensitive to adenylate levels. This occurs through the TSC-Rheb circuit, independent of amino acid sensing pathways (Hoxhaj et al., 2017). AMPK and mTORC1 are major drivers of metabolic reprogramming with differential roles in Teff and Treg cells (Bantug et al., 2018; Buck et al., 2015). Therefore, AICAR accumulation and associated reduction in mTORC1 activity may contribute to impaired generation of MTHFD2i-treated Th1 cells and enhanced generation and lineage stabilization observed in Treg cells. It is noted that there is also a reciprocal relationship between mTORC1 and MTHFD2 as mTORC1 can drive accumulation of the transcription factor ATF4, which in turn can induce MTHFD2 expression (Ben-Sahra et al., 2016).

Spatiotemporal compartmentalization of MTHFD2 may also contribute to the effects of MTHFD2-deficiency. Specifically, *de novo* purine biosynthesis occurs in dynamic multi-enzyme complexes called purinosomes that allows for tight regulation of metabolic flux (Pedley and Benkovic, 2017). Purinosomes colocalize with mitochondria and are closely tied to mitochondrial function and metabolism. Moreover, this link is mediated by mTOR activity (French et al., 2016). Given that GAR, SAICAR, and AICAR accumulation occur in Th1 and Treg cells with transient drug exposure and purine depletion occurs only in Th17 cells with extended exposure, these metabolons may be partly mediating the differential effects of MTHFD2i on CD4 T cell subsets.

Metabolic flux is tied to epigenetic regulation in that many metabolites can serve as cofactors in epigenetic modifications (Sharma and Rando, 2017). In addition to nucleotide synthesis, 1C metabolism is critical for generating the universal methyl-donor SAM for protein and DNA methylation. Transfer of the methyl group from SAM yields SAH, and SAM/SAH ratios can affect the function of key methyltransferases (Mentch et al., 2015). At homeostatic conditions, differential SAM/SAH ratios may dictate distinct global methylation levels in CD4 T cell subsets. In T cells, carbon tracing experiments show that 1C units derived from glucose-derived and exogenous serine do not contribute to the methionine cycle (Ma et al., 2017). Thus, methylation changes observed with MTHFD2i may not be a direct consequence of reduced formate pools. Our finding that Th17 cells treated with MTHFD2i have altered methylation patterns may indicate a disruption of methylation directly as well as indirect effects of impaired differentiation.

MTHFD2 has also been reported to play non-enzymatic roles that may contribute to the pro-inflammatory actions of this enzyme. In cancer cells, MTHFD2 can have nuclear functions, co-localizing with DNA replication sites to promote cell cycle progression (Sheppard et al., 2015). In murine stem cells, MTHFD2 was shown to modulate DNA repair to maintain genomic stability (Yue et al., 2020). In renal cell carcinoma, MTHFD2 was found to be crucial for metabolic reprogramming via mRNA methylation (Green et al., 2019). While these roles for MTHFD2 are not mutually exclusive with our findings, the ability of formate to rescue many of the phenotypes of MTHFD2 deficiency in T cells suggest that MTHFD2 enzymatic activity is the primary driver of T cell proliferation and fate.

As a drug target, MTHFD2 has been largely considered in anti-cancer settings. It may also provide a promising anti-inflammatory target and offer fewer adverse effects compared to currently available anti-folates given low expression in most adult tissues. For instance, one target of methotrexate, DHFR, is expressed extensively in adult tissues (Nilsson et al., 2014). As such, methotrexate can be associated with a variety of adverse effects including gastrointestinal toxicities that lead to treatment withdrawal. Moreover, despite its extensive history, the mechanism of action of methotrexate remains poorly understood (Cronstein and Aune, 2020). The highly regulated expression of MTHFD2 and potential for redundancy with the cytosolic MTHFD1 pathway may result in selective dependencies of specific cell populations on MTHFD2. Together our findings identify MTHFD2 as a critical metabolic checkpoint in CD4 T cells. While there is broad overlap between cancer cell biology and the biology of rapidly proliferating T cells, we find that CD4 subsets display an additional sensitivity to mitochondrial 1C metabolism in determining cell fate and function. It is likely that targeting MTHFD2 in cancer therapies may restrain anti-tumor immunity. However, settings such as colorectal carcinoma, where Th17-mediated inflammation contributes to disease, may be well suited to MTHFD2 inhibitors. This T cell sensitivity may make MTHFD2 inhibitors an effective form of immunotherapy in settings of CD4 T-cell driven inflammation beyond cancer, and lead to fewer adverse effects than currently available therapeutics.

## AUTHOR CONTRIBUTIONS

A.S., G.A., and J.C.R. designed research; A.S., G.A., K.V., D.R.H., K.L.B., M.M.W., D.L.G.S.K.S., S.N.F., A.M.C., P.F., X.X., and J.C.G.C. performed research; A.S., G.A., X.Y., A.K.M., J.D.R., and J.C.R. analyzed data; and A.S. and J.C.R. wrote the paper.

## ACKNOWLEDGEMENTS

We thank members of the Rathmell lab for contributing to this project. We thank the Mangalam Lab (University of Iowa) for performing the EAE experiment, Thomas Aune for RNAseq data, and Sitryx Therapeutics Limited (Oxford, UK) for constructive input and performing the DTH experiment. We thank J. Cools (VIB) for providing the pMx-U6-gRNA-GFP construct. We thank Nello Mainolfi, Vipin Suri, Adam Friedman, and Mark Manfredi from Raze Therapeutics, Inc. (Boston, MA) for providing the Raze 1459 compound. This work was supported by R01s DK105550, HL136664, and AI153167 (J.C.R.), AI137075 (A.K.M.), T32 DK101003 (K.V.), and T32 GM007347 (A.S.).

## METHODS

### Key Resources Table

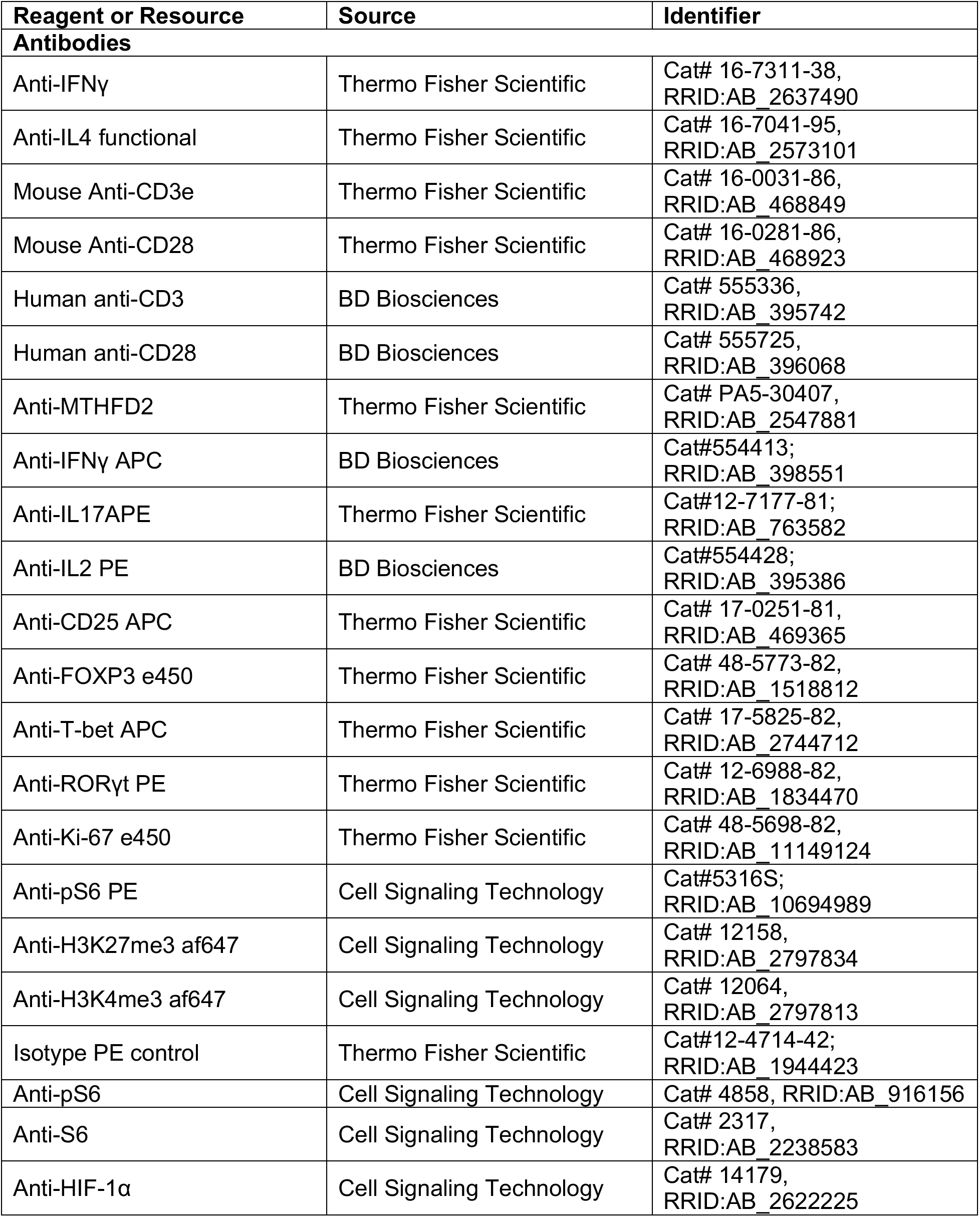
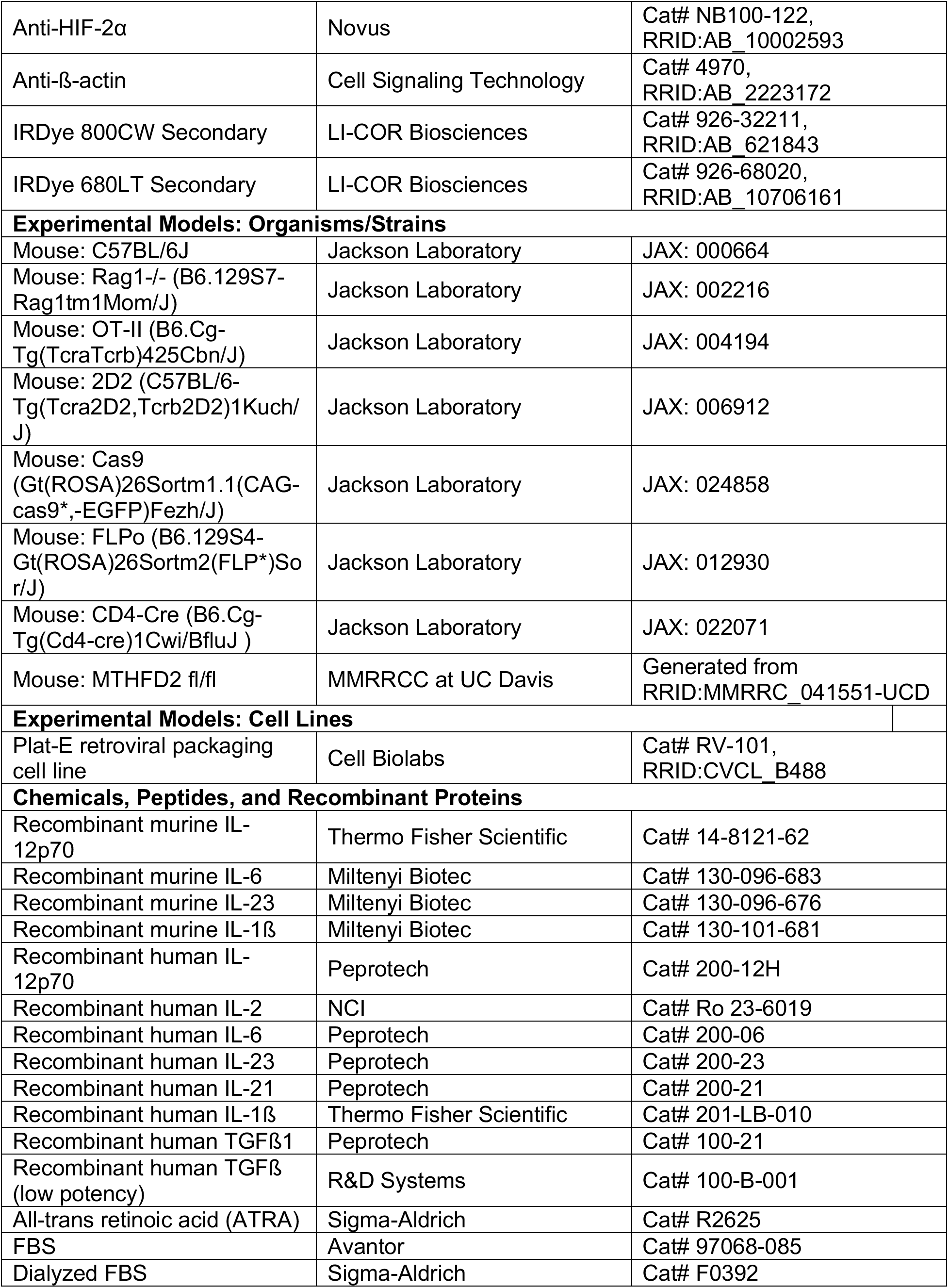
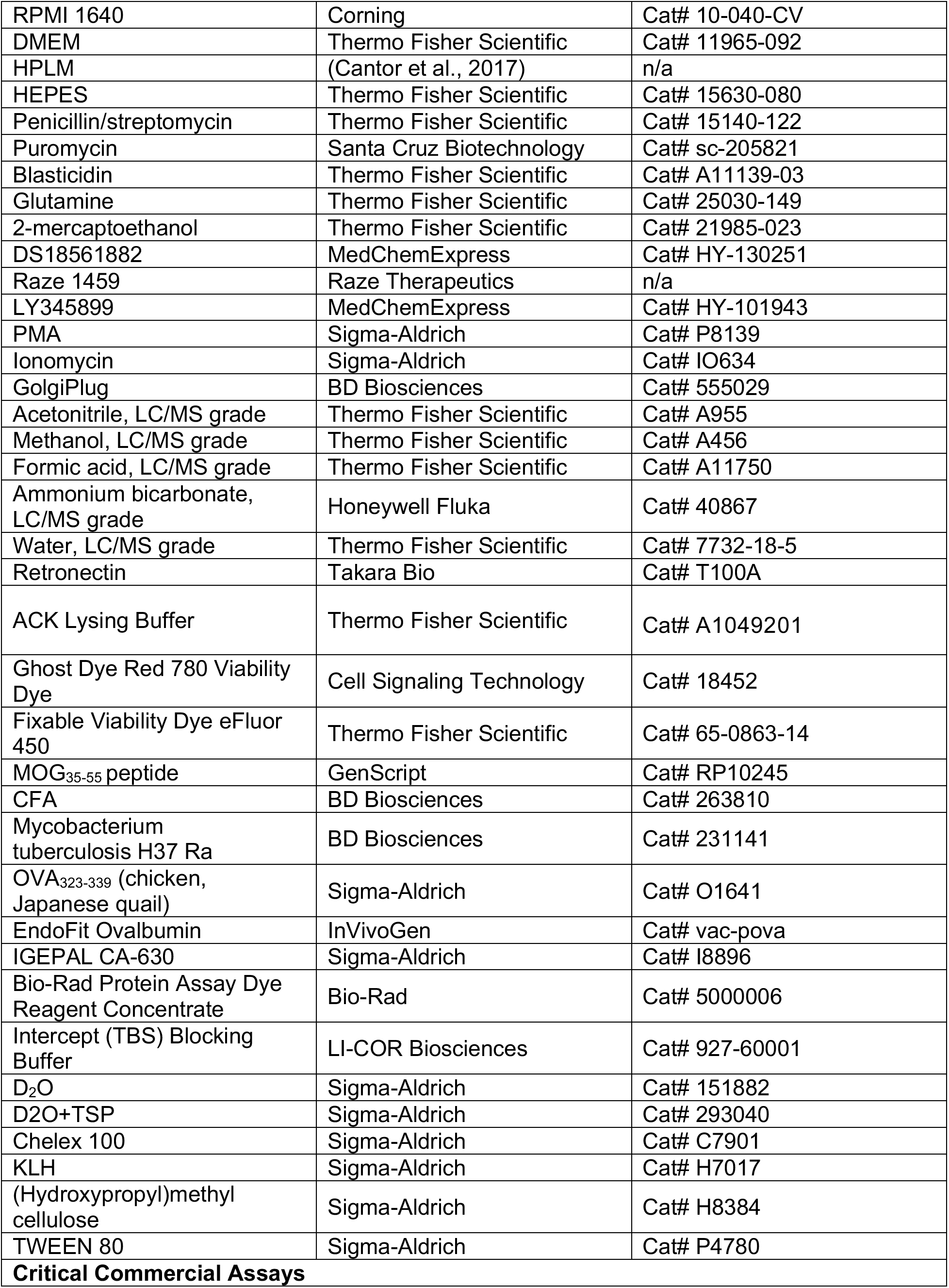
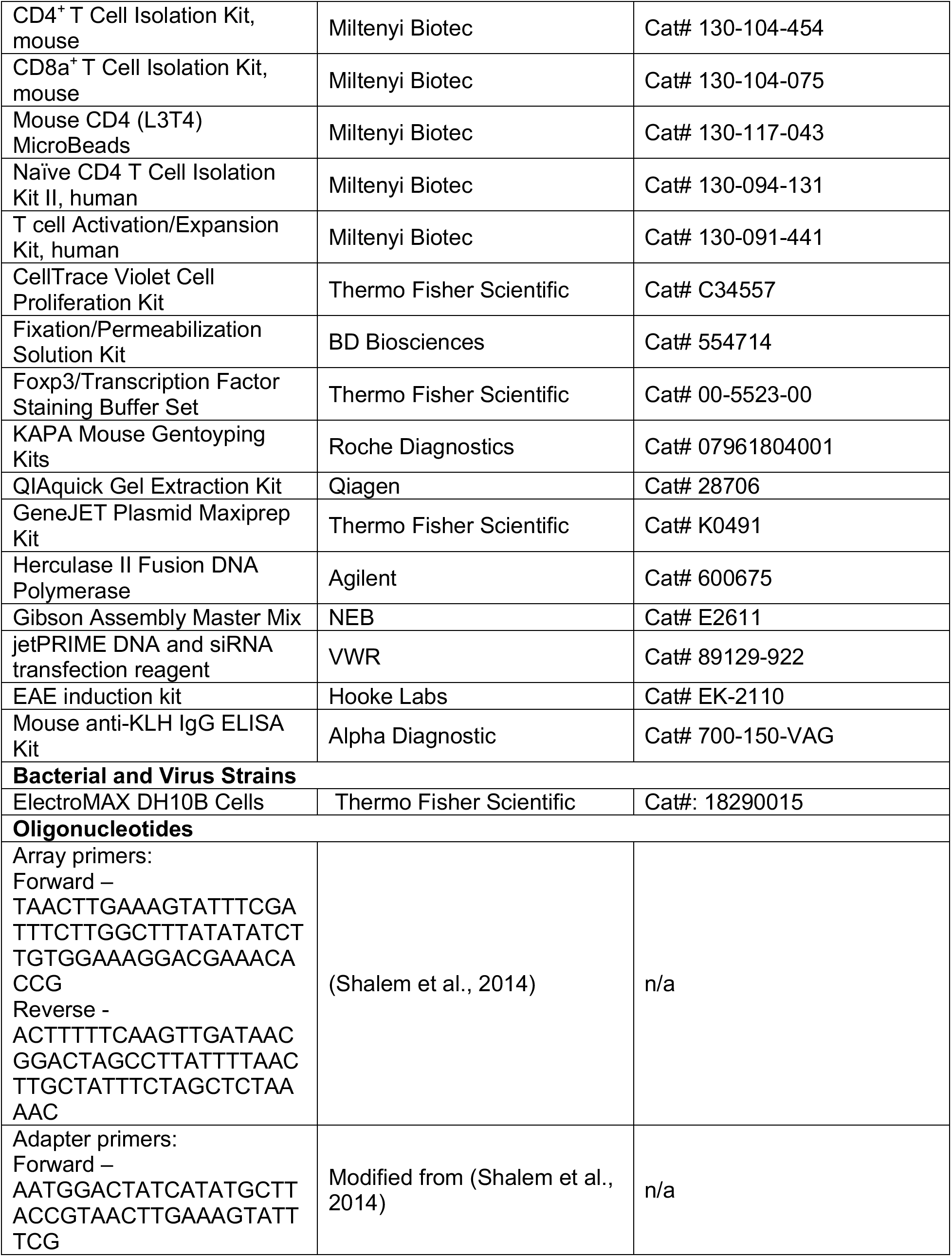
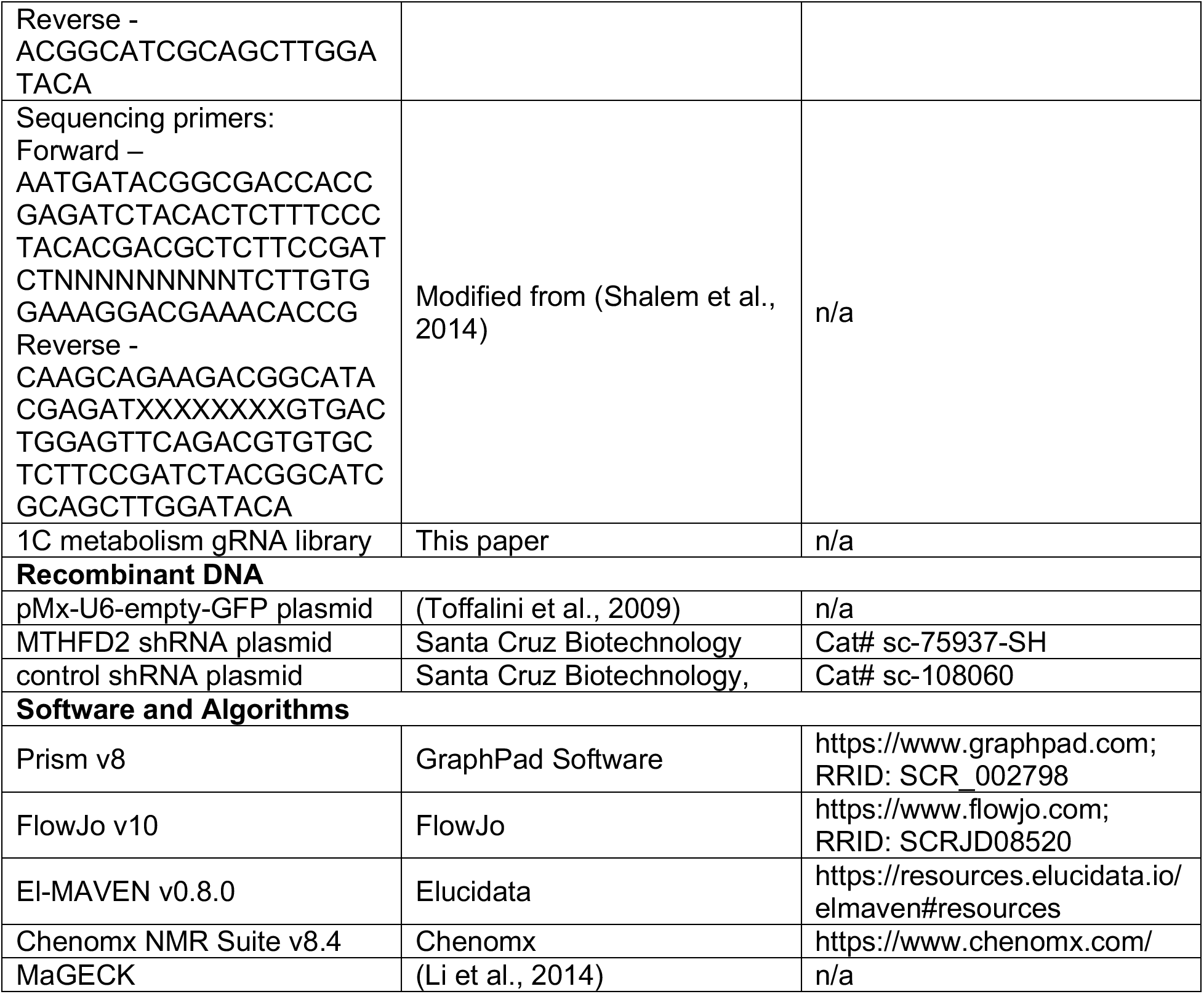

### Lead Contact and Materials Availability

Further information and requests for resources and reagents should be directed to and will be fulfilled by the Lead Contact, Dr. Jeffrey Rathmell.

### Experimental Model and Subject Details

#### Mice

All experiments were performed at Vanderbilt University, University of Iowa, or Fidelta animal facility in accordance with Institutional Animal Care and Utilization Committee (IACUC)-approved protocols and conformed to all relevant regulatory standards. Mice were housed in pathogen-free facilities in ventilated cages with *ad libitum* food and water and at most 5 animals per cage. Eight-to sixteen-week-old male and female mice were used for all animal experiments. All mice except MTHFD2^fl/fl^ mice described below were obtained from Jackson Laboratory and were treatment-naive until the start of study.

OT-II mice were crossed to Cas9 mice to generate OT-II/Cas9 double-transgenic strain. Animals were genotyped for transgenic T-cell receptor (TCR) and Cas9 allele. MTHFD2^fl/fl^ mice were generated from C57BL/6N-Atm1Brd Mthfd2tm1a(EUCOMM)Wtsi/BayMmucd by Mutant Mouse Resource & Research Centers (MMRRC) at UC Davis. These mice were first crossed to FLPo mice. The F1 mice were then crossed to CD4-Cre mice for several generations to obtain a strain with MTHFD2^fl/fl^ CD4-Cre^+^ conditional knockout (KO) and WT littermates. Animals were genotyped for floxed and Cre alleles.

#### Cell Lines

Plat-E retroviral packaging cell line was maintained at 37°C with 5% CO_2_ in DMEM media supplemented with 10% FBS, 100U/mL penicillin/streptomycin, 1μg/mL puromycin, and 10μg/mL blasticidin to maintain expression of viral packaging genes.

### Method Details

#### In vitro mouse CD4 T cell activation and differentiation

Primary murine CD4 T cells were isolated from the spleens and lymph nodes of mice using a negative isolation kit (Miltenyi Biotec) according to the manufacturer’s instructions. The cells were cultured at 37°C with 5% CO_2_ in RPMI 1640 media supplemented with 10mM HEPES, 50μM 2-mercaptoethanol, 100U/mL pennicilin/streptomycin, and 2mM glutamine, unless otherwise stated. Primary CD4 T cells were activated using plate-bound anti-CD3 (3μg/mL) and anti-CD28 (5μg/mL) antibodies at 125,000 cells/well for 96-well plate and 1 million cells/well for 24-well plate. Cells were cultured for 3 to 4 days as specified with subset-specific cytokines and antibodies to promote differentiation - Th1: IL-12p70 (10ng/mL), IL-2 (100U/mL), anti-IL-4 (10μg/mL), anti-IFNγ (1μg/mL); Th17: IL-6 (50ng/mL), TGFβ (1ng/mL), IL-23 (10ng/mL), IL-1b (10ng/mL), anti-IL4 (10μg/mL), anti-IFNγ (10μg/mL); Treg: TGFβ (1.5ng/mL), IL-2 (100U/mL), anti-IL4 (10μg/mL), anti-IFNγ (10μg/mL). Low potency TGFβ (R&D Systems) was used instead for Treg differentiation as indicated. MTHFD2 inhibitors were dosed at 100nM and 500nM for DS18561882 and 2.5μM for Raze 1459. For intracellular and transcription factor stains, cells were first stained with viability dye, fixed and permeated using appropriate kits, then stained for proteins. For cytokine stains, cells were stimulated for four hours with 12-myristate 13-acetate (PMA) and ionomycin in the presence of GolgiPlug for four hours, then processed as other intracellular stains. For MTHFD2 staining, cells were fixed in Fixation/Permeabilization Solution (BD Sciences) followed by ice-cold methanol.

#### Th17 suppression assay

Primary CD4 T cells were activated and differentiated to Th17 cells for 72 hours with 0nM, 100nM, or 500nM DS18561882. Th17 cells were then collected, washed, and re-seeded into fresh complete media with responder CD8 T cells at equal numbers on anti-CD3/anti-CD28 coated plates. Responder CD8 T cells were isolated from the spleens and lymph nodes of mice using a negative isolation kit (Miltenyi Biotec) and stained with 5μM CellTrace Violet (CTV) cell proliferation dye. CTV dilution was assessed by flow cytometry after 72 hours of co-culture.

#### Treg stability assay

Primary CD4 T cells were activated and differentiated to Treg cells for 72 hours. Treg cells were then collected, washed, and re-seeded into fresh complete media supplemented with IL-2 (10ng/mL) and either DMSO control or MTHFD2i (2.5μM Raze 1459 for mouse experiment, 2μM DS18561882 for human experiment). Cells were cultured for an additional 48 hours before analysis.

#### In vitro human CD4 T cell culture conditions

PBMCs were collected under an approved IRB from anonymous healthy blood donors. Naïve CD4 T cells were isolated using the Naïve CD4 T Cell Isolation Kit II (Miltenyi Biotec) according to the manufacturer’s instructions. All human T cells were cultured at 37°C with 5% CO_2_ in human plasma-like medium (HPLM) (Cantor et al., 2017) supplemented with 10% dialyzed FBS. In MTHFD2i experiments, T cells were activated in the presence of either DMSO control or 2μM DS18561882. Th1 cultures were activated with CD2/3/28 activation beads (Human T cell Activation/Expansion kit, Miltenyi Biotec) at a 2:1 ratio of cells to beads with anti-IL-4 (2μg/mL), IL-2 (100U/mL), and IL-12p70 (5ng/mL). Th17 cultures were activated with non-tissue culture treated plates with plate-bound anti-CD3 (3μg/mL), anti-CD28 (1μg/mL), and anti-ICOS (1μg/mL) antibodies. Media was supplemented with anti-IL-4 (2μg/mL), anti-IFNγ (2μg/mL), IL-23 (50ng/mL), IL-1β (50ng/mL), TGFβ1 (5ng/mL), IL-21 (25ng/mL) and IL-6 (40ng/mL). iTreg cultures were activated similarly with plate-bound anti-CD3 (4μg/mL) and anti-CD28 (3μg/mL) antibodies in media supplemented with anti-IL-4 (2μg/mL), anti-IFNγ (2μg/mL), TGFβ1 (5ng/mL), IL-2 (100U/mL), and all-trans retinoic acid (ATRA, 10nM). Th1 and iTreg cultures were also supplemented with IL-2 from day 3.

For shRNA knockdown experiments, cells were pulsed with either an MTHFD2 shRNA or a control shRNA plasmid using the Invitrogen Neon Transfection System (1 pulse at 2,100V, 20ms). Knockdown efficiencies were determined by flow cytometry staining of MTHFD2 after 3 days.

#### Immunoblotting

Cells were lysed with base lysis buffer containing 1% IGEPAL CA-630, 200mM NaCl, and 50mM Tris pH 8.0 on ice for 30 minutes. The base lysis buffer was supplemented with the protease inhibitors aprotinin (5ug/mL), leupeptin (5ug/mL), sodium fluoride (0.9mM), dithiothreitol (DTT, 1mM), sodium vanadate (1mM), and beta-glycerophosphate (20mM).

Lysates were centrifuged for 15 minutes at 4°C to recover supernatant and quantified for protein concentration using Protein Assay Dye Reagent Concentrate (Bio-Rad). 40ug of protein was loaded per well for polyacrylamide gel electrophoresis using Mini-PROTEAN Precast Polyacrylamide Gels (Bio-Rad). Western blotting was performed using low fluorescence PVDF membrane (Bio-Rad). Transfer was accomplished using 1X Towbin Transfer Buffer Containing 20% methanol at 300mA for 1 hour. Blots were blocked for 1 hour using Intercept (TBS) Blocking Buffer (LI-COR Biosciences), before incubation with primary antibody overnight at 4°C.

Blots were incubated for 1hr at room temperature with IRDye Secondary antibodies (LI-COR Biosciences) and were visualized by near infrared fluorescence via Li-COR Odyssey CLx imager. The antibodies used for westerns were: B-actin (1:1000), Phospho-S6 Ribosomal Protein (Ser235/236) (1:2000), S6 Ribosomal Protein (1:1000), HIF-1α (1:1000), and HIF-2α (1:1000).

#### Mass spectrometry (MS) metabolomics

Cultured cells were collected and washed with PBS. Each sample was resuspended in precooled extraction buffer (by volume, 40% methanol, 40% acetonitrile, 20% water, and 0.5% formic acid) at the ratio of 75μL of buffer per 1 million cells. The samples were then neutralized by 15% NH_4_HCO_3_ aqueous solution at 8.8 vol% of the extraction buffer. Cell extracts were stored in −80°C freezers prior to analysis. Samples were thawed on ice, and insoluble contents were pelleted by centrifugation (16,000xg, 10 min) at 4°C. To further precipitate proteins, 40μL of supernatant was diluted in 40μL of methanol, followed by incubation on dry ice for 1 hour. Then the samples were spun down (16,000xg for 10 min) at 4°C and supernatants were transferred into vials and stored at 4°C until analysis.

Samples were analyzed by Q Exactive Plus Quadrupole-Orbitrap Mass Spectrometer (Thermo Scientific) coupled with hydrophilic interaction chromatography (HILIC). 10μL of each sample was injected into the LC system. Metabolites were separated by a XBridge BEH Amide Column (2.5μM, 150mm × 2.1mm, Waters) with a 25 min gradient from Solvent B (acetonitrile) to Solvent A (95 vol% H_2_O and 5 vol% acetonitrile solution with 20 mM NH_4_OAc, 20 mM NH_3_·H_2_O, pH 9.4). LC-MS method was reported in Anal. Chem. 2019, 91, 3, 1838–1846 (Wang et al., 2019). Raw data were converted into .mzXML format, and data analysis was performed using El-MAVEN (v0.8.0).

#### Proton nuclear magnetic resonance spectroscopy (^1^H-MRS)

Conditioned culture media and cells were collected following 72 hours of culture and processed as previously reported (Govindaraju et al., 2000). Briefly, for quantification of metabolites from conditioned supernatant, a total of 50 μL D2O and 50 μL of 0.75% sodium 3-trimethylsilyl-2,2,3,3-tetradeuteropropionate (TSP) in D2O was added to 500 μL media and transferred to 5-mm NMR tubes (Wilmad-LabGlass, Kingsport, TN). For quantification of intracellular metabolites, a total of 180 μL of a master mix of D2O and of 0.75% TSP in D2O were added to dried soluble cell extract and transferred to 3-mm NMR tubes (Wilmad-LabGlass, Kingsport, TN). ^1^H-MRS spectra were acquired on an Avance III 600 MHz spectrometer equipped with a Triple Resonance CryoProbe (TCI) (Bruker) at 298 K with 7500-Hz spectral width, 32,768 time domain points, 32 scans (supernatant) or 256 scans (cell extracts), and a relaxation delay of 2.7 seconds. The water resonance was suppressed by a gated irradiation centered on the water frequency. The spectra were phased, baseline corrected, and referenced to TSP using Chenomx NMR Suite. Spectral assignments were based on literature values.

#### CRISPR screening

1C metabolism gRNA library was curated by referencing the Mouse CRISPR Knockout Pooled Library (Brie) (Addgene, Pooled Library #73632) (Doench et al., 2016). Four gRNA sequences for each gene and ten non-targeting controls, flanked by the following adapter sequences were purchased as an oligo pool from Twist Bioscience: GGAAAGGACGAAACACCGXXXXXXXXXXXXXXXXXXXXGTTTTAGAGCTAGAAATAGCAAG TTAAAATAAGGC. The library was further prepared for transduction following published methods (Anderson et al., 2017; Shalem et al., 2014) with several modifications. Briefly, additional sequences were attached by PCR using Array primers and Herculase II Fusion DNA Polymerase. After purification by gel extraction using QIAquick Gel Extraction Kit, the fragment was cloned into the retroviral expression vector pMx-U6-gRNA-GFP using Gibson Assembly Master Mix. The resultant plasmid pool was amplified by electroporation into ElectroMAX DH10B Cells and plated on ampicillin plates to obtain enough colonies for 50-fold coverage of the library. DNA was isolated using GeneJET Plasmid Maxiprep Kit.

DNA was transfected into Plat-E retroviral packaging cell line using Polyplus jetPRIME DNA and siRNA transfection reagent. Media was changed 24 hours post transfection, and viral supernatant was collected after an additional 48 hours of culture. Meanwhile, CD4 T cells were isolated from the spleen and lymph nodes of OT-II/Cas9 double-transgenic mice and activated with splenocytes irradiated at 30 Gy and OVA323-339 peptide (10μg/mL). 48 hours post T cell activation, the viral supernatant was spun onto retronectin treated non-tissue culture plates at 2000xg for 2 hours at 32°C. Activated T cells were transferred to the plates and spun for an additional 15 minutes, then replaced in the incubator. On day 3 post T cell activation, a sample of the cells were collected at 1000-fold representation of the library. Additionally, 3 million live GFP^+^ cells were adoptively transferred into each Rag1-/- mice by tail vein injection. On days 1, 3, 5, and 7 post adoptive transfer, mice were sensitized with intranasal ovalbumin protein. On day 8 post adoptive transfer, lungs and spleens were collected, and T cells isolated by positive selection using CD4 (L3T4) Microbeads.

Genomic DNA from cells were extracted with Kapa Express Extract Kit. gRNA sequences were amplified by two rounds of PCR with two technical replicates: first round with adapter primers and second round with barcoded Illumina sequencing primers. The amplicons were then purified by gel extraction, combined at equimolar, and sequenced on the Illumina NovaSeq platform. FASTQ files were input in MAGeCK (Li et al., 2014) for statistical analysis.

#### In vivo EAE for immunohistochemistry and flow cytometry

2 separate methods of EAE induction were used. In the first method, female C57BL/6 mice aged 8-12 weeks were injected subcutaneously with 0.2mL myelin oligodendrocyte glycoprotein (MOG) peptide in Complete Freund’s Adjuvant (CFA) emulsion and i.p. 100ng pertussis toxin (PTX) (Hooke Labs). The PTX injection was repeated 24 hours later. In the second method, CD4 T cells were isolated from 2D2 mice aged 8-12 weeks. Cells were activated with splenocytes irradiated at 30Gy and MOG_35-55_ peptide (10μg/mL) and Th17 cytokines. On day 4 post activation, cells were transferred to fresh media with IL-23 (20ng/mL) to rest for an additional 3 days. On day 7 post activation, cells were re-stimulated with plate-bound anti-CD3 and anti-CD28 antibodies. 48 hours post restimulation, 10 million live cells were resuspended in 100μl PBS with 2% FBS and injected into Rag1^-/-^ mice by tail vein injection. Mice were observed daily for clinical severity.

Once mice started exhibiting paralysis of both hind limbs, they were sacrificed for further analysis. For IHC, the entire spinal column was dissected out and fixed in 1%PFA. For T cell isolation, the spinal cord was dissected out and digested in 300U/mL collagenase and 50U/mL DNase at 37 °C for 45 min. The digested cord was filtered through a 70μm filter and spun on a 18.6%/62.4% Percoll gradient at 2400rpm for 30min at room temperature. Immune cells were collected from the middle layer, washed, and stained for flow cytometry.

#### In vivo MTHFD2i in EAE

Female C57BL/6 mice aged 8-12 weeks were purchased from Jackson Laboratories and maintained at the University of Iowa animal facility in accordance with NIH and institutional guidelines. EAE was induced in mice by subcutaneous immunization in both flanks with 100μg of MOG_35-55_ peptides (MEVGWYRSPFSRVVHLYRNGK) emulsified in CFA containing Mycobacterium tuberculosis H37Ra (100μg/mouse). PTX (100 ng) was administered i.p. at days 0 and 2 post immunization (Shahi et al., 2019). Animals were divided into two groups and treated i.p with 10mg/kg MTHFD2i LY345899 or DMSO/PBS vehicle control every day starting day 4 post EAE induction till the end of the experiment. Mice were observed daily for clinical severity and scored according to the following scale: 0-no clinical disease, 1-loss of tail tonicity, 2-hind limb weakness, 3-partial hind limb paralysis, 4-complete hind limb paralysis, and 5-moribund/death. Mice were euthanized at score of 4 for analysis.

#### In vivo MTHFD2i in DTH

Female BALB/c mice aged 8 weeks were purchased from Charles River and maintained at Fidelta animal facility in accordance with AAALAC and institutional guidelines. KLH-DTH was induced by subcutaneous immunization on day 0 with 100μL of 4mg/mL KLH and 0.5mg/mL Complete Freund’s Adjuvant emulsion. Mice were challenged on day 7 in the pinna of the left ear with 4mg/mL KLH antigen dissolved in 0.9% NaCl solution, while the right ear pinna was challenged with 0.9% NaCl solution. Animals were treated from days 0-10 twice daily (BID) via the per oral route (PO) with 10 mL/kg vehicle (5% DMSO and 95% (0.5%) HPMC/(0.1%) Tween80 in distilled water), or DS18561882 at a dose of 100 and 300 mg/kg and 10 mL/kg. Body weight was measured daily from days 0-10, ear thickness was measured on day 7 prior to challenge and then on days 8-10. On day 10 terminal blood was collected by carotid artery bleeding. Serum was prepared from coagulated blood samples by centrifugation at 3500rpm/15min, separated and stored at −80 °C for subsequent KLH-IgG analysis by ELISA. ELISAs were performed in accordance with the manufacturer’s instructions. Postmortem, a circular piece of both ear pinna, 8mm in diameter, was cut out with a biopsy punch, weighed on a precise analytical balance and recorded.

### Statistical analysis

Statistical analyses were performed with Prism software (v8). Statistically significant results are labelled (* p < 0.05, ** p < 0.01, *** p < 0.001, **** p ≤ 0.001). Error bars show mean ± standard deviation unless otherwise indicated. All experiments were carried out in triplicate biological replicates unless otherwise stated. FACs plots shown are representative of n = 3 biological replicates.

