## Supplemental figures and legends for "MTHFD2 is a Metabolic Checkpoint Controlling Effector and Regulatory T Cell Fate and Function"

#### Supplemental Figure Legends

##### Supplementary Figure 1: 1C metabolism and MTHFD2 expression in CD4 T cells

(A) Growth rate by protein mass in CD4 T cells activated on anti-CD3/anti-CD28 and cultured in indicated cytokine conditions over 72 hours.

(B) Glucose (left), lactate (middle), and glutamine (right) uptake by cells from (A) at 72 hours post activation corrected for cell proliferation by protein mass (one-way ANOVA).

(C) Change in gRNA abundance from *in vivo* CRISPR/Cas9-based screening of 1C metabolism genes in primary CD4 T cells in a lung inflammation model. Ranked by depletion score (analyzed by MaGECK).

##### Supplementary Figure 2: Potential mechanisms mediating MTHFD2i effects

(A) Intracellular adenosine nucleotide levels in Th1, Th17, and Treg cells treated with either vehicle or 500nM DS18561882 for 72 hours (unpaired t-test).

(B) Total cellular ROS (DCFDA, left) and mitochondrial superoxide (mitoSOX; right) levels 72 hours post activation with anti-CD3/anti-CD28 in Th1, Th17, and Treg cells treated with 0, 100, or 500nM DS18561882 (one-way ANOVA).

(C) Fold change in SAM (left) and SAH (middle) levels in Th1, Th17, and Treg cells five days post activation compared to resting cells (one-way ANOVA). SAM/SAH ratio (right) in Th1, Th17, and Treg cells (one-way ANOVA).

(D) H3K4-trimethylation levels in Th1, Th17, and Treg cells treated with 0nM, 100nM, or 500nM DS18561882 measured by flow cytometry (one-way ANOVA).

(E) Immunoblot of HIF-2α in Th17 cells treated with vehicle or 500nM DS18561882.

##### Supplementary Figure 3: Effect of MTHFD2i on CD4 T cells infiltrating spinal cord in EAE

(A-B) Frequency (top) and total count (bottom) of FoxP3+ (A) and IFNγ$^+$ (B) CD4 T cells in the spinal cord of mice immunized with $MOG_{35-55}$/CFA and PTX and injected i.p. daily with vehicle or LY345899 at 10mg/kg (unpaired t-test).

**Supplementary Figure 4: Effect of MTHFD2i in DTH**

(A) Percent change in body weight from day 0 to 10 in mice immunized with KLH/CFA on day 0 and challenged with KLH on day 7, treated twice daily with PO vehicle, 100, or 300mg/kg DS18561882 (mean±SEM).

(B) Change in ear thickness from day 7 to 10 in the same mice as (A) (mean±SEM).

### Supplemental Figure 1

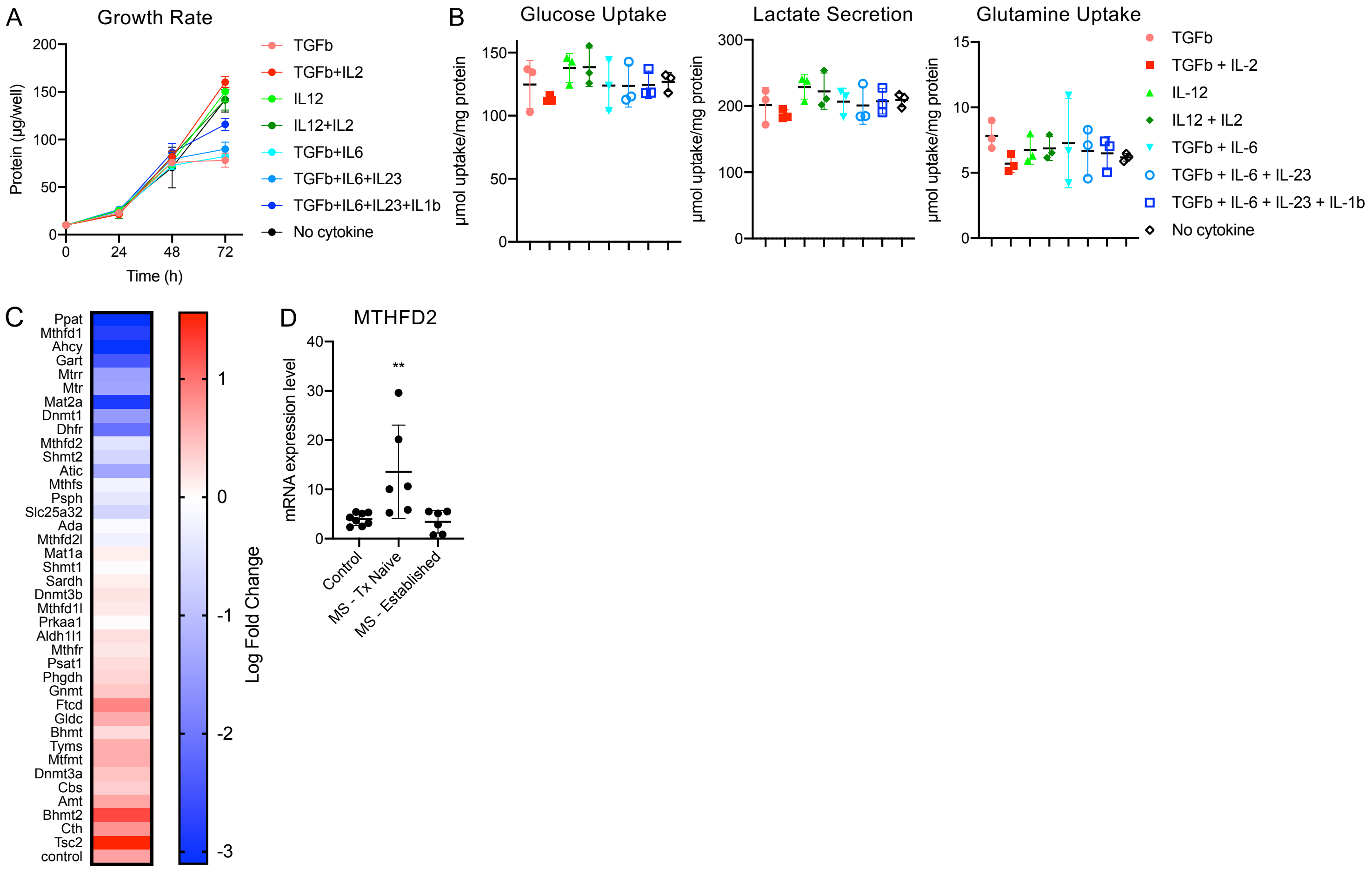

### Supplemental Figure 2

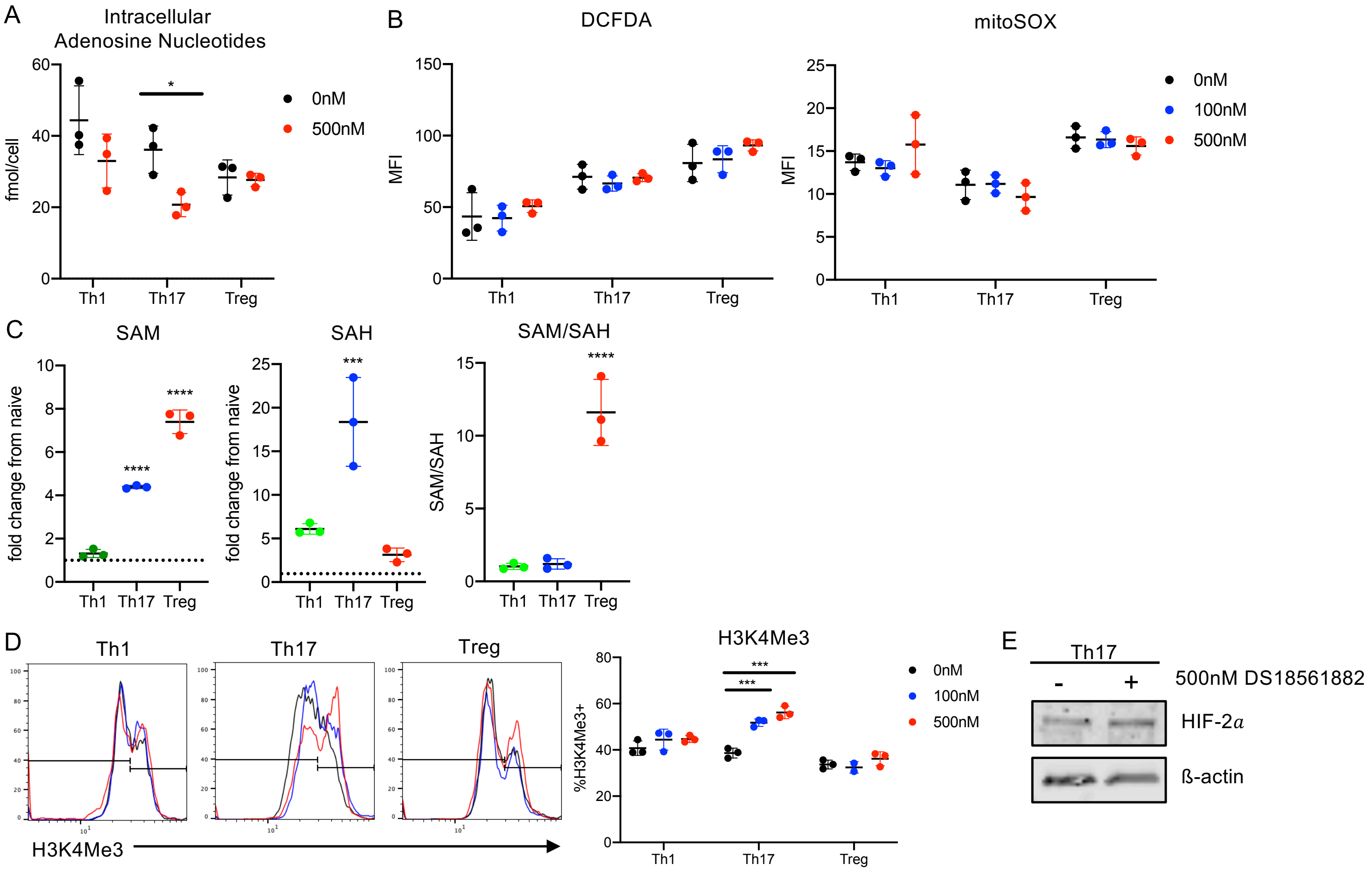

### Supplemental Figure 3

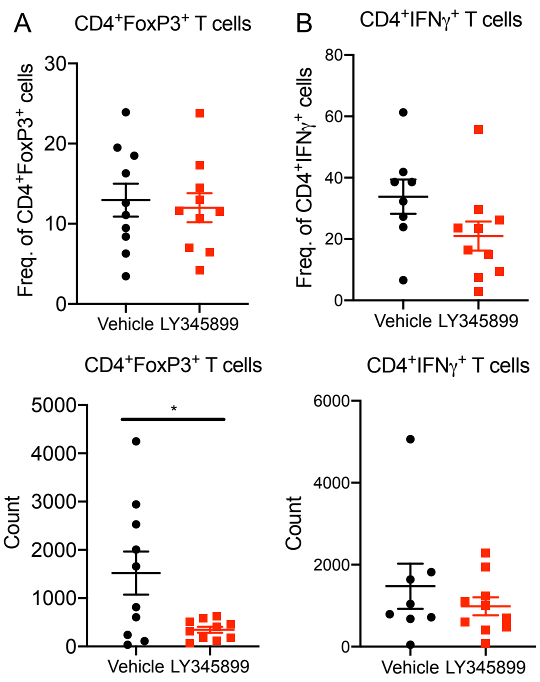

### Supplemental Figure 4

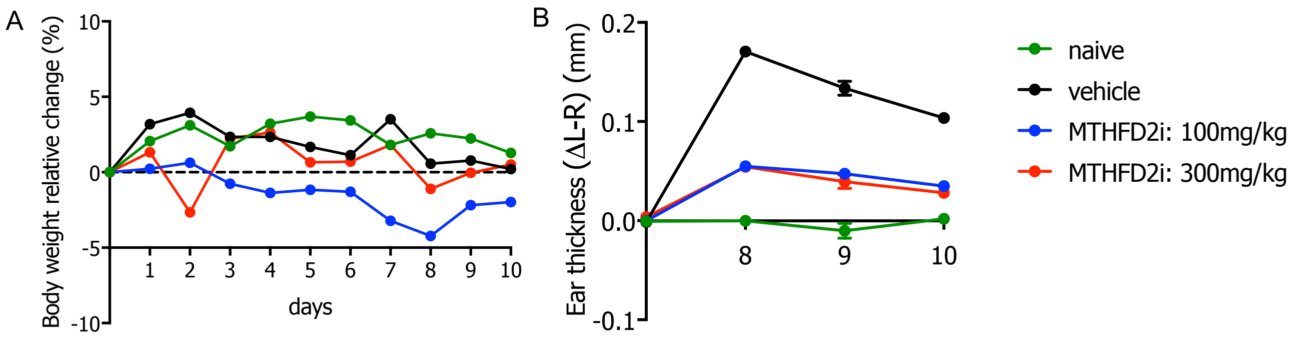
